# Eye position signals in the dorsal pulvinar during fixation and goal-directed saccades

**DOI:** 10.1101/681130

**Authors:** Lukas Schneider, Adan-Ulises Dominguez-Vargas, Lydia Gibson, Igor Kagan, Melanie Wilke

**Affiliations:** Decision and Awareness Group, Cognitive Neuroscience Laboratory, German Primate Center, Leibniz Institute for Primate Research, Kellnerweg 4, Goettingen, 37077, Germany; Department of Cognitive Neurology, University of Goettingen, Robert-Koch-Str. 40, Goettingen, 37075, Germany; Leibniz ScienceCampus Primate Cognition, Kellnerweg 4, Goettingen, 37077, Germany

**Author notes:** Corresponding author: Igor Kagan, German Primate Center, Leibniz Institute for Primate Research, Kellnerweg 4, Goettingen, 37077, Germany. Shared last authorship (Igor Kagan, Melanie Wilke).

**Keywords:** memory saccades, eye movements, orbital gaze position, reference frames, macaque

## Abstract

Most sensorimotor cortical areas contain eye position information thought to ensure perceptual stability across saccades and underlie spatial transformations supporting goal-directed actions. One pathway by which eye position signals could be relayed to and across cortical areas is via the dorsal pulvinar. Several studies demonstrated saccade-related activity in the dorsal pulvinar and we have recently shown that many neurons exhibit post-saccadic spatial preference long after the saccade execution. In addition, dorsal pulvinar lesions lead to gaze-holding deficits expressed as nystagmus or ipsilesional gaze bias, prompting us to investigate the effects of eye position. We tested three starting eye positions (−15°/0°/15°) in monkeys performing a visually-cued memory saccade task. We found two main types of gaze dependence. First, ∼50% of neurons showed an effect of static gaze direction during initial and post-saccadic fixation. Eccentric gaze preference was more common than straight ahead. Some of these neurons were not visually-responsive and might be primarily signaling the position of the eyes in the orbit, or coding foveal targets in a head/body/world-centered reference frame. Second, many neurons showed a combination of eye-centered and gaze-dependent modulation of visual, memory and saccadic responses to a peripheral target. A small subset showed effects consistent with eye position-dependent gain modulation. Analysis of reference frames across task epochs from visual cue to post-saccadic target fixation indicated a transition from predominantly eye-centered encoding to representation of final gaze or foveated locations in non-retinocentric coordinates. These results show that dorsal pulvinar neurons carry information about eye position, which could contribute to steady gaze during postural changes and to reference frame transformations for visually-guided eye and limb movements.

**New & Noteworthy:** Work on the pulvinar focused on eye-centered visuospatial representations, but position of the eyes in the orbit is also an important factor that needs to be taken into account during spatial orienting and goal-directed reaching. Here we show that dorsal pulvinar neurons are influenced by eye position. Gaze direction modulated ongoing firing during stable fixation, as well as visual and saccade responses to peripheral targets, suggesting involvement of the dorsal pulvinar in spatial coordinate transformations.

## Introduction

Information about eye position is ubiquitous in the primate brain and is critical for visually-guided behavior. Neurons modulated by the position of the eyes in the orbit have been reported in brain stem nuclei (Luschei and Fuchs, 1972; Hernández et al., 2019), superior colliculus (Van Opstal et al., 1995; Campos et al., 2006), inferior colliculus (Porter et al., 2006), thalamus (Schlag-Rey and Schlag, 1984; Wyder et al., 2003; Tanaka, 2007), cerebellum (Noda and Warabi, 1982), and numerous cortical regions including visual and fronto-parietal cortices (Bizzi, 1968; Andersen et al., 1990; Squatrito and Maioli, 1996; Wang et al., 2007; Morris et al., 2013). These eye position signals might subserve different functions. In the visual/oculomotor domain eye position signals could enable stable vision, the discrimination between self- and external motion, precise stimulus localization across eye movements, and post-saccadic updating (Sommer and Wurtz, 2008; Wurtz et al., 2011a). During inter-saccadic periods, these signals might enable stable fixation, including vergence of the eyes and smooth pursuit (Squatrito and Maioli, 1996). In the context of visually-guided limb and body movements, eye position signals might serve to transform the retinal location of visual stimuli into head-, limb- or trunk-centered reference frames (Andersen et al., 1993; Colby, 1998; Pouget and Snyder, 2000). In most experimental situations, the head is immobilized, so the position of eyes in the orbit is equivalent to gaze direction, or angle.

Taking the current gaze angle into account is also important for the visual guidance of movements since visual inputs enter the brain in eye-centered (retinocentric) coordinates, but control of limb movements requires locating the objects in respect to the body and the world. A common model of how the brain deals with those spatial reference frame transformations are the so called “gain fields”, manifested as a modulation of sensory-evoked, motor preparation or movement-evoked responses by the current eye position on a single neuron level, such that the neural population response simultaneously represents the retinal and body-centered stimulus location (Andersen et al., 1990; Pouget and Snyder, 2000; Salinas and Abbott, 2001; Cohen and Andersen, 2002). Neurons that exhibit a modulation of visual responses by eye position were first reported in the intralaminar nuclei of the thalamus (Schlag et al., 1980) and the superior colliculus (SC) in cats (Peck et al., 1980), and later in monkeys in the ventral (retinotopically-organized) pulvinar (Robinson et al., 1990). A systematic assessment of spatial tuning at different eye positions revealed gain field properties across numerous cortical regions, including parietal areas (e.g. lateral intraparietal area LIP, ventral intraparietal area VIP, medial intraparietal area MIP, area 7a), the frontal eye fields (FEF), posterior cingulate and middle superior temporal area MST (Andersen et al., 1990; Galletti et al., 1995; Squatrito and Maioli, 1996; Bremmer et al., 1999, 2002; Dean and Platt, 2006; Lehky et al., 2016; Caruso et al., 2018).

It is not entirely clear how this eye position information is distributed across cortical areas. A region that could possibly fulfill this function is the dorsal pulvinar (Sherman and Guillery, 2002; Arcaro et al., 2018). The dorsal pulvinar (dPul), consisting of the medial pulvinar and dorsal part of the lateral pulvinar, is in a good anatomical and functional position to transfer gaze-related information to and across fronto-parietal and superior temporal cortices (Grieve et al., 2000; Wurtz et al., 2011a; Bridge et al., 2015; Halassa and Kastner, 2017). The dorsal pulvinar receives direct input from the intermediate and deep layers of the superior colliculus (Benevento and Standage, 1983; Baldwin and Bourne, 2017) and is reciprocally interconnected with prefrontal (FEF, dorsal lateral prefrontal cortex dlPFC), dorsal premotor (PMd), posterior parietal (LIP, MIP, VIP, area 7) and superior temporal sulcus regions such as MST and temporal parietal occipital area TPO (Romanski et al., 1997; Gutierrez et al., 2000; Cappe et al., 2012; Yeterian and Pandya, 1989; Kaas and Lyon, 2007). Beyond its anatomical connections, electrophysiological and lesion studies suggest a critical role of the dorsal pulvinar in spatial attention (Petersen et al., 1987; Fiebelkorn et al., 2019) as well as visuomotor processes including the control of eye movements. Specifically, response properties of dorsal pulvinar neurons in monkeys partially resemble its diverse cortical projection targets such as parietal cortex, e.g. enhancement for visual stimuli that indicate an upcoming saccade target (Robinson, 1993). While the dorsal pulvinar is not retinotopically organized and its neurons have large receptive fields, they discharge in the context of saccade tasks in visual cue and saccade execution phases, exhibiting overall preference for contralateral visual cue, peri- and post-saccadic responses (Petersen et al., 1985; Benevento and Port, 1995; Dominguez-Vargas et al., 2017).

Unilateral pharmacological inactivation of the dorsal pulvinar results in decreased ability to shift attention into the contralesional field (Robinson and Petersen, 1992) and a saccade choice bias towards the ipsilesional field (Wilke et al., 2010, 2013). Human patient studies are largely consistent with those results, showing contralesional spatial deficits (Karnath et al., 2002; Arend et al., 2008; Van der Stigchel et al., 2010). At the same time, inactivation or structural lesions have comparatively modest effects on basic saccade parameters, mostly consisting of decreased latencies for ipsilesional visually-guided saccades and increased latencies for contralesional memory-guided saccades (Wilke et al., 2010, 2013). In contrast to primary oculomotor regions (e.g. SC, FEF or internal medullary lamina (IML)/central thalamus (Schiller and Tehovnik, 2005; Tanaka and Kunimatsu, 2011)), relatively high currents (>150 µA) are necessary to evoke saccades and those are small, infrequent and might depend on behavioral context (Dominguez-Vargas et al., 2017). Apart from eye movement selection, the dorsal pulvinar also plays a critical role in other visuomotor behaviors such as reaching and grasping; its neural activity correlates with reach movements (Cudeiro et al., 1989; Acuna et al., 1990) and inactivation/lesions in monkeys and humans lead to hand- and space-specific deficits in reach and grasp tasks (Wilke et al., 2010, 2018).

Based on those anatomical and functional results it has been proposed that the dorsal pulvinar is involved in reference frame transformations critical for eye-hand behavior, although this has not been directly demonstrated (Grieve et al., 2000; Bridge et al., 2015). In fact, very basic questions such as whether dorsal pulvinar neurons carry eye position signals and how those interact with visual cue responses have not been addressed. Even the effect of static eye position on ongoing firing has not been tested, although the presence of tonic firing during initial and post-saccadic fixation (Dominguez-Vargas et al., 2017) as well as nystagmus and smooth pursuit deficits following dorsal pulvinar inactivation/lesions suggest that it might be involved in gaze stabilization as well (Ohtsuka et al., 1991; Wilke et al., 2010, 2018).

Here we investigated the effect of eye position on initial fixation, visual, memory, saccade and post-saccadic responses in monkeys performing a memory-guided saccade task. We demonstrate that about half of dorsal pulvinar cells are modulated by steady gaze position before visual cue onset and long after the saccade. We also demonstrate a gaze-dependent modulation of visual and saccadic activity, providing a possible substrate for head- or body-centered spatial representations.

## Materials and Methods

All experimental procedures were conducted in accordance with the European Directive 2010/63/EU, the corresponding German law governing animal welfare, and German Primate Center institutional guidelines. The procedures were approved by the responsible government agency (LAVES, Oldenburg, Germany).

### Animal preparation

Two adult male rhesus monkeys (*Macaca mulatta*) C and L weighing 8 and 9 kg respectively, were used. In an initial surgery, monkeys were implanted with a magnetic resonance imaging (MRI) compatible polyetheretherketone (PEEK) headpost embedded in a bone cement headcap (Palacos with Gentamicin, BioMet, USA) anchored by ceramic screws (Rogue Research, Canada), under general anesthesia and aseptic conditions. MR-visible markers were embedded in the headcap to aid the planning of the chamber in stereotaxic space with the MR-guided stereotaxic navigation software Planner (Ohayon and Tsao, 2012). A separate surgery was performed to implant a PEEK MRI-compatible chamber(s) (inside diameter 22 mm) allowing access to the pulvinar (Monkey C, right hemisphere: center at 0.5A / 14.5L mm, tilted −11P / 27L degrees; Monkey L, right hemisphere: center at −3.12P / 20.2L mm, tilted: −18P/37L degrees; Monkey L, left hemisphere: center at −3P/20L, tilted: - 18P/-38L). After confirming chamber positioning with a post-surgical MRI, a partial craniotomy was made inside the chamber. The exposed dura was covered with a silicone elastomer (Kwik-sil, World Precision Instruments, USA) to reduce the granulation tissue growth and dura thickening.

### MRI imaging

Monkeys were scanned in a 3T MRI scanner (Siemens Magnetom TIM Trio). Full-head T1-weighted scans (3D magnetization-prepared rapid gradient-echo, MPRAGE, 0.5 mm isometric) were acquired before and after chamber implantation, in awake (monkey C) or anaesthetized (monkey L) state, using either built-in gradient body transmit coil and custom single loop receive coil, or custom single loop transmit and 4-channel receive coil (Windmiller Kolster Scientific, USA).

In addition to pre- and post-implantation scans, similar T1-weighted scans as well as T2-weighted (rapid acquisition with relaxation enhancement, RARE, 0.25 mm in plane, 1 mm slice thickness) scans were periodically acquired during the course of experiments, either in awake (monkey C) or sedated (monkey L) state, to confirm electrode positioning. T1- and T2-weighted scans were co-registered and transformed into “chamber normal” (aligned to the chamber vertical axis) and to anterior commissure – posterior commissure (AC-PC) space for electrode targeting and visualization. These images were acquired with the chamber and the grid filled with gadolinium (Magnevist, Bayer, Germany)/saline solution (proportion 1:200), with tungsten rods inserted in predefined grid locations, for alignment purposes.

### Gaze modulation task

Monkeys sat in a dark room in custom-made primate chairs with their heads restrained 30 cm away from a 27’’ LED display (60 Hz refresh rate, model HN274H, Acer Inc. USA), covering a horizontal range of 100 visual degrees. The gaze position of the right eye was monitored at 220 Hz using an MCU02 ViewPoint infrared eyetracker (Arrington Research Inc. USA). All stimulus presentation and behavioral control tasks were programmed in MATLAB (The MathWorks, Inc. USA) and the Psychophysics Toolbox (Brainard, 1997).

The structure of the task is shown in Figure 1A. A trial started with the onset of the fixation spot of 1° diameter either at the center or 15 ° left or right to the center of the screen. After the monkey acquired and held fixation within a 5° radius for 500 ms, a peripheral cue (also 1° diameter) was displayed for 300 ms signaling the upcoming saccade target location. For each trial, one out of eight cue/target locations were used. These eight positions were arranged in a virtual rectangular around the initial fixation spot, at 0°, 15° or −15° horizontally and 0°, 10° or −10° vertically, resulting in 24 different spatial conditions (eight target locations for each of the three initial fixation positions). Monkeys were required to maintain fixation throughout the cue period and also throughout the subsequent memory period (1000 ms), after which the central fixation spot disappeared, allowing monkeys to saccade to the instructed target location. This time point will be referred to as the “Go signal”. After a saccade and fixation inside a 5° radius window surrounding the remembered target location for 200 ms the target became visible. After additional 500 ms of peripheral fixation the trial was completed and the monkey obtained a liquid reward after a delay of 200 ms. The inter-trial interval for successful and unsuccessful trials was 2500 ms. All initial fixation positions (3) and retinocentric target locations (8) were pseudo-randomized.

**Figure 1.**
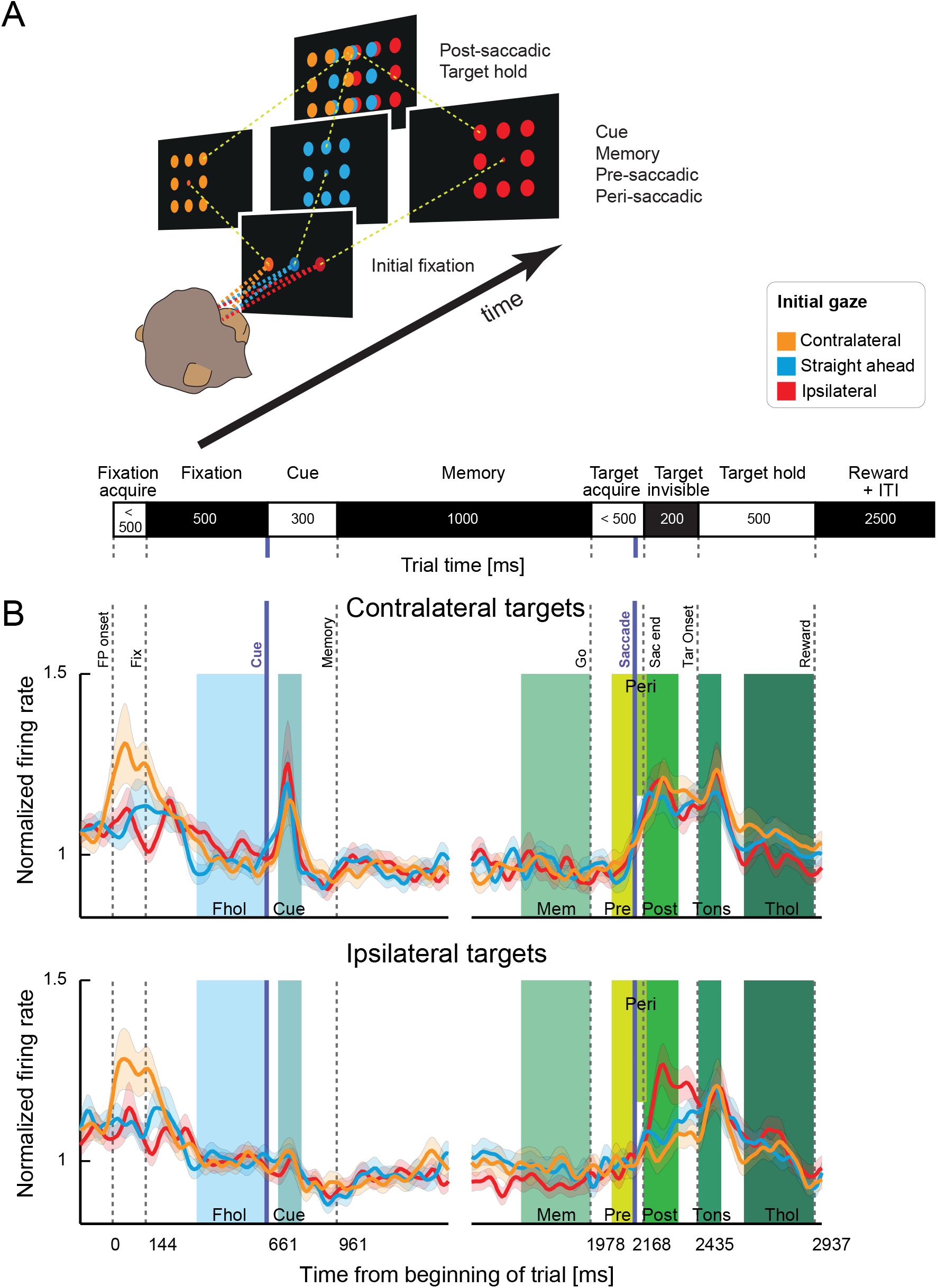
Task and population average. **A:** Task conditions. A trial started at one out of three initial fixation positions (Fixation). One out of eight target positions surrounding the fixation spot was flashed (Cue). Monkeys had to remember the location while keeping fixation (Memory), until the fixation spot disappeared, prompting a saccade to the target (Target acquire). After the monkey held fixation for a short period (Target invisible), the target reappeared as a confirmation (Target hold). **B:** Normalized population PSTHs and standard errors across units for each initial gaze position for contralateral and ipsilateral targets (relative to initial gaze position). Vertical lines indicate average onset of events across all trials: fixation spot onset (FP onset), the monkey acquiring fixation (Fix), the cue onset, the cue offset and beginning of the memory period, the offset of the central fixation point (Go), the saccade onset, the monkey acquiring the invisible target location (Sac end), the onset of the confirmation target (Tar Onset), and the end of the trial (Reward). Discontinuous traces indicate two different alignments to cue onset and saccade onset (purple lines). Colored areas mark analysis epochs: fixation hold (Fhol), cue onset (Cue), memory (Mem), pre-saccadic (Pre), peri-saccadic (Peri), post-saccadic (Post), target onset (Tons), and target hold (Thol) (see Materials and Methods).

## Data analysis

### Saccade definition

Saccade velocity was calculated sample by sample as the square root of the sum of squared interpolated (220 Hz to 1 kHz) and smoothed (12 ms moving average rectangular window) horizontal and vertical eye position traces, and then smoothed again (12 ms moving average rectangular window). Saccade onset was defined as the first eye position change that exceeded a starting velocity threshold of 300°/s.

### Dataset and unit selection criteria

All recorded voltage drops that surpassed an online visually determined threshold were defined as potential spikes. Spike sorting was done in Offline Sorter v.3.3.5 (Plexon, USA), using a waveform template algorithm after defining templates by manually clustering in principle component space.

In total, 325 single and multi-units were recorded in the dorsal pulvinar in 22 sessions where monkeys performed the gaze modulation task (monkey C; right hemisphere: 134, monkey L; left hemisphere: 191). Out of these, 275 units (93 monkey C, 182 monkey L) fulfilled analysis selection criteria (stable discriminability across time and reasonable signal to noise ratio – assessed by inspection). 268 out of these 275 units (86 monkey C, 182 monkey L) were recorded for at least 4 successful trials for each of the initial gaze positions. These 268 units were used for initial gaze analysis. 238 out of these 268 units (60 monkey C, 178 monkey L) were recorded for at least 4 successful trials for each combination of initial gaze and retinocentric target location. These 238 units were used for further gaze dependent analysis.

### Epoch definitions and modulation

For each trial, and each epoch of interest, firing rates were computed by counting spikes within the epoch and dividing by the epoch duration. The following epochs were analyzed: fixation hold (last 300 ms of central fixation), cue onset (50 ms to 150 ms after cue onset), memory (last 300 ms of the memory period), pre-saccadic (100 to 10 ms before saccade onset), peri-saccadic (10 ms before to 50 ms after saccade onset), post-saccadic (first 150 ms after acquiring the invisible peripheral target), target onset (20 ms to 120 ms after the target became visible), and target hold (last 300 ms of fixating the peripheral target).

For analysis of basic response types in the entire population (**Supplemental Figure S1**), trials were either grouped by retinocentric or screen-centered target location. We performed two ANOVAs on firing rates of each unit in each epoch: a two-way ANOVA with factors initial gaze position and retinocentric target location and a one-way ANOVA dependent on screen-centered target location. Additionally we computed retinocentric hemifield preferences for all units that showed a main effect of retinocentric target location in the respective epoch and screen-centered hemifield preferences for all units that showed an effect of screen-centered target location. To this end, data from all contralateral and ipsilateral hemifield targets were combined. Hemifield tuning in each epoch was determined by unpaired t-tests comparing firing rates in ipsilateral trials to firing rates in contralateral trials. The hemifield with the higher firing rate was marked, if there was a significant difference.

Enhancement or suppression of neuronal activity in each epoch was defined by paired t-tests comparing firing rates to a respective preceding baseline epoch, independently for ipsilateral and contralateral trials. For the fixation hold epoch, inter-trial interval served as baseline, while for cue onset and memory epochs the fixation hold epoch served as baseline. The memory epoch served as baseline for all subsequent epochs. Enhancement or suppression was reported, if either ipsilateral, contralateral, or both types of trials showed significant difference to fixation baseline. In rare cases where one hemifield would show a significant enhancement, while the other hemifield showed suppression, the unit was reported to have a bidirectional response.

### Peri-stimulus time histograms (PSTHs)

Spike density functions were computed at a bin size of 10 ms using a Gaussian kernel (σ=20 ms). Example PSTHs show spike densities averaged across all trials with the same initial fixation position. For population PSTHs, we first normalized spike density functions for each unit by dividing by the average firing rate in the fixation hold epoch (across all trials), and computed average spike density and standard error (across units) for each initial gaze position and contralateral/ipsilateral retinocentric target positions individually.

### Gaze position dependent analysis

To evaluate effects of gaze position on neuronal activity of each unit, we analyzed initial and final gaze position effects. An effect of initial gaze position was determined by a one-way ANOVA on firing rates during the fixation hold epoch with 3 initial gaze positions as the independent factor, and an effect of final gaze position was determined by a one-way ANOVA on firing rates during the target hold epoch with 15 final gaze positions as the independent factor. To dissociate horizontal and vertical gaze tuning, an additional two-way ANOVA was performed with factors horizontal and vertical final gaze position.

To evaluate if gaze dependence varied systematically with horizontal and/or vertical gaze position, units with a main effect along the respective dimension were further grouped by monotonic and non-monotonic gaze-dependent activity. Monotonic gaze dependence was defined by a) a significant difference between firing rates for the most peripheral positions (unpaired t-test) and b) no significant opposite difference in firing rates between any two neighboring positions. All other units that showed an effect in the ANOVA were classified as non-monotonic. Units showing non-monotonic initial gaze preference were further grouped into central preference (highest activity for straight ahead gaze) and peripheral preference (lowest activity for straight ahead gaze). For better visualization, firing rates for each unit were normalized by dividing average responses for each of the three initial/final gaze positions by the maximum average firing rate across all positions.

To see how task contingency affected gaze dependent responses, we compared initial (fixation hold) and final (target hold) gaze contralaterality indices across all units. Contralaterality indices were computed as (C-I)/(C+I), where C and I are the average firing rates for contralateral and ipsilateral gaze positions, relative to the center of the screen. For final gaze contralaterality indices 6 ipsilateral and 6 contralateral gaze positions were combined. Straight ahead gaze positions were not used for this analysis.

### Gaze-dependent modulation of spatially-contingent task epochs

To evaluate the presence of not purely retinocentric response fields for each unit, average firing rates in each epoch were tested with a two-way ANOVA, using the factors initial gaze position and retinocentric target location.

To evaluate the relationship of gaze and cue tuning across units, we first correlated initial gaze and retinocentric cue spatial preferences and second tested if there was a difference in response modulation by comparing absolute strength of gaze and cue spatial preferences using Wilcoxon’s signed rank test. In total, we performed four different comparisons: spatial preferences were either taken as raw firing rate differences (C-I, where C and I are the average firing rates in contralateral and ipsilateral trials) or contralaterality indices (C-I)/(C+I); and gaze preference was either derived from fixation hold (Fhol) or Cue onset (Cue) epoch. Spatial cue preference was computed using all 3 gaze positions, combining all cues contralateral or ipsilateral relative to the current gaze.

### Modulation of retinocentric encoding by gaze position

To evaluate the presence of gain fields for each unit and each epoch, we used a model-free approach that does not assume any specific shape of the tuning function. First, we computed confidence intervals for tuning vector length and direction, independently for each initial gaze position using hierarchical bootstrapping. 1000 bootstrapping iterations were performed for each unit, epoch and initial gaze position, sampling 10 trials (with replacement) for each retinocentric target position. Normalized tuning vectors for each sampled trial were computed with direction equal to the direction of the retinocentric target position and the length equal to the firing rate. The average tuning vector for the current bootstrap iteration was then computed as the sum of normalized firing rate vectors across all sampled trials. To evaluate significance of gain and tuning vector shifts, the 95% confidence intervals of differences in amplitude and direction of the bootstrapped tuning vectors for each pair of initial gaze positions was computed, independently for each epoch. Units in which at least for one of these comparisons between gaze positions, the confidence interval of amplitude or direction differences did not overlap with zero were marked as showing gain field properties or response field shifts, respectively. This analysis was performed only on units that showed a main effect of retinocentric target location in the respective epoch in the two-way ANOVA mentioned before.

To estimate gain effect size, we calculated the percent change in the amplitude of the average bootstrapped tuning vector across three initial gaze positions for each unit as 100*(Amax-Amin)/Amax, where Amax is the amplitude of the largest and Amin the amplitude of the smallest of the three average tuning vectors.

### Reference frame estimation

To see if pulvinar responses were better explained by a retinocentric or screen-centered reference frames (i.e. relative to the locations on the screen, which could signify head-, body-, or world-centered representation, since the head was immobilized relative to the screen and the body orientation was not explicitly controlled), we grouped the trials either by retinocentric target location or by the target location on the screen, and compared average correlation coefficients (ACCs) in both arrangements for each unit (Mullette-Gillman et al., 2005). For each reference frame, ACCs were computed by correlating responses for two out of three initial gaze positions at a time, and then averaging the three correlation coefficients. For screen-centered encoding, we used only locations that were available for both initial gaze positions for every correlation pair. To estimate significance, we performed 1000 bootstrap iterations using 80% of the trials for each location at a time. This allowed deriving 95% confidence intervals for both retinocentric and screen-centered ACCs. Significant encoding in the respective reference frame was reported if a) the confidence interval was above zero, b) the confidence interval for this reference frame was above the ACC for the other reference frame, and c) the ACC for this reference frame was above the confidence interval for the other reference frame. In other words, both confidence intervals should be above or below the unity line diagonal.

## Results

We recorded single- and multi-unit activity in two monkeys performing a memory-guided saccade task with variation of initial gaze (fixation) position (Figure 1A). Monkeys had to fixate at one of three initial positions, hold fixation while a spatial cue was presented at one of 8 target locations and after a memory period make a saccade towards the cued target location. These 8 potential target locations were arranged in a rectangle around the initial fixation spot, with the same spatial arrangement relative to the fixation spot for each of the three initial gaze positions, see Figure 1A and Materials and Methods. Since the monkeys were head-fixed in the straight-ahead direction, the position of the eyes in the orbit is equivalent to gaze direction.

A total of 268 units in the dorsal pulvinar were studied in two monkeys (monkey C: 86, monkey L: 182). For most analyses beyond the effects during initial fixation we focused on the units with more than 4 trials for each combination of initial gaze and target position (N=238, monkey C: 60 monkey L: 178), see Materials and Methods. Normalized population PSTHs of these 238 units are displayed in Figure 1B. In the population average we found no apparent dependence on the initial gaze position in the fixation hold epoch, an enhanced contralateral cue response and post-saccadic enhancement.

Neurons in this sample exhibited diverse activity patterns; a summary of response modulation (enhancement, suppression, or bidirectional modulation) and spatial preferences across task epochs is given in **Supplemental Figure S1**. In short, our sample contained 48% of units with initial fixation responses (24% enhanced, 24% suppressed, relative to pre-trial ‘initialize trial epoch’), 31% with spatial cue dependence (11% contralateral preference, 7% ipsilateral preference, and 13% with a main effect of target location without hemifield preference), 44% with memory delay period activity (13% enhanced, 31% suppressed), 19% with pre-saccadic activity (8% enhanced, 11% suppressed), and 67% with post-saccadic activity (47% enhanced, 19% suppressed). A detailed analysis of dPul neuronal response properties in a more extensive non-overlapping sample will be given in a separate paper (see also (Dominguez-Vargas et al., 2017)). Here, we focus on the activity patterns pertaining to different positions of the eyes in the orbit.

### Gaze-dependent activity during initial fixation

A substantial portion of units (128 out of 268, 48%) were significantly modulated by the gaze position before the cue was presented (one-way ANOVA). Many of these units maintained gaze dependence after cue presentation (47 out of 128, 37%). Three example units shown in Figure 2 illustrate tonic responses during the initial fixation and subsequent trial epochs. Focusing first on the initial fixation epoch, top example (Figure 2A) shows contralateral gaze preference and two other examples (Figure 2B,C) show ipsilateral preference. Our first question was if such gaze-dependent activity typically increased or decreased towards peripheral contra- or ipsilateral gaze positions, or if more units prefer the straight ahead direction, as has been demonstrated in the visual cortex (Durand et al., 2010; Przybyszewski et al., 2014). To this end, we classified the units which showed a main effect of initial gaze position into units with monotonic gaze-dependent effects (showing peripheral preference), and units that showed either central or peripheral non-monotonic gaze dependence (Materials and Methods).

**Figure 2.**
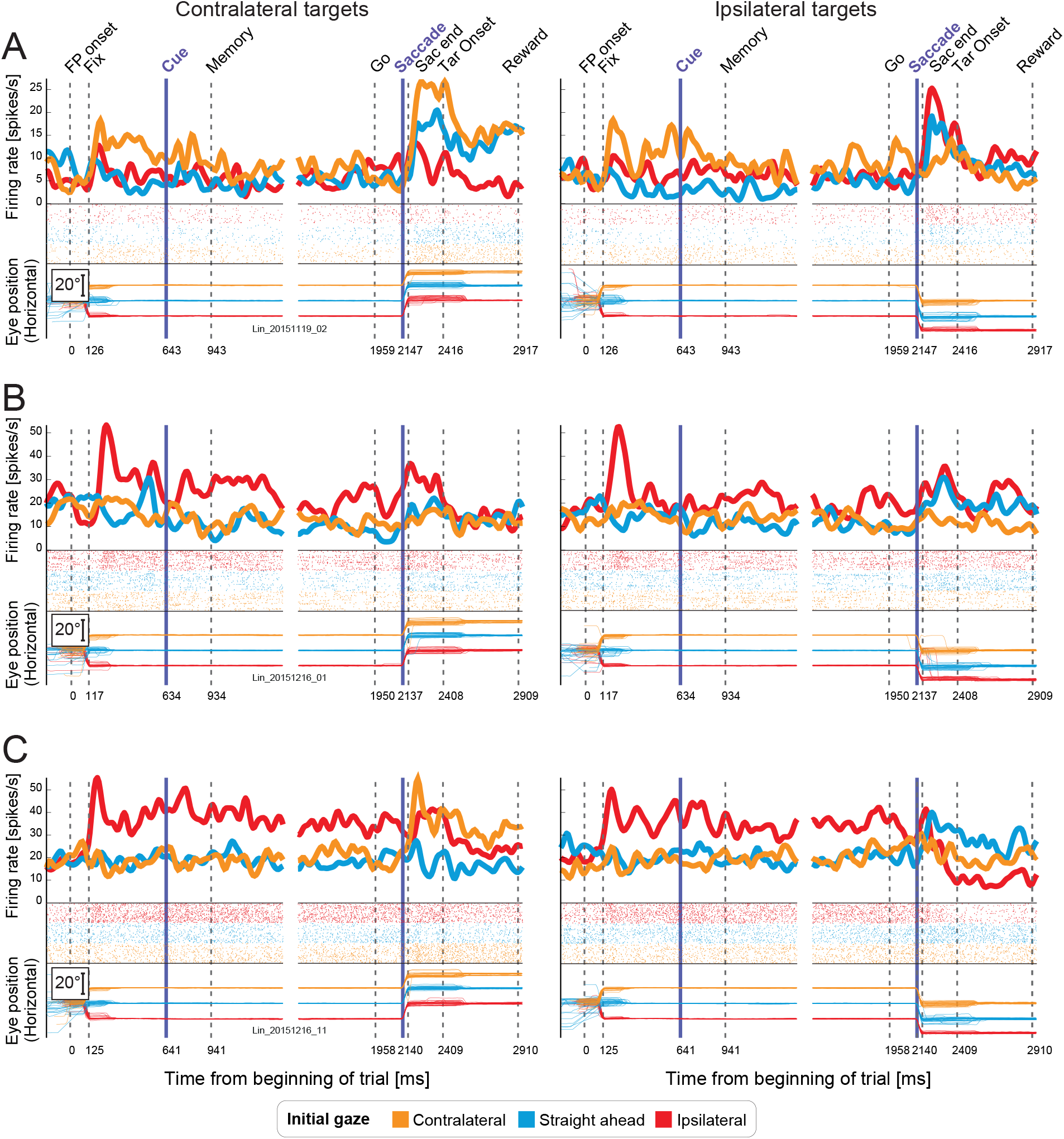
Example units with tonic gaze-dependent activity. Raster plots, resulting spike density functions, and horizontal eye traces separately for each initial fixation position, for contralateral (left) and ipsilateral targets (right) relative to initial gaze position. Colors depict the three initial gaze positions (orange: contralateral, blue: straight ahead, red: ipsilateral). **A:** Peripheral contralateral gaze preference, **B:** and **C:** Peripheral ipsilateral gaze preference.

Figure 3A shows normalized firing rates during fixation hold for each unit and each of the three initial gaze positions, grouped by the monotonicity and gaze direction preference. Monotonic responses were more frequent than non-monotonic (91/128 and 37/128 units), but contralateral and ipsilateral preferences were almost balanced in monotonic units (41 and 50 units respectively). Furthermore, in non-monotonic units there was no bias towards the straight ahead direction (18 vs. 19 units). If anything, we found an overall preference for peripheral positions (110 units: all monotonic and non-monotonic peripheral-preferring vs. 18 central-preferring units). MR-guided reconstruction of recording sites revealed that units with significant gaze dependence were distributed throughout the sampled locations (mostly in the dorsal medial pulvinar), without a systematic clustering of gaze-dependent patterns (Figure 3B).

**Figure 3.**
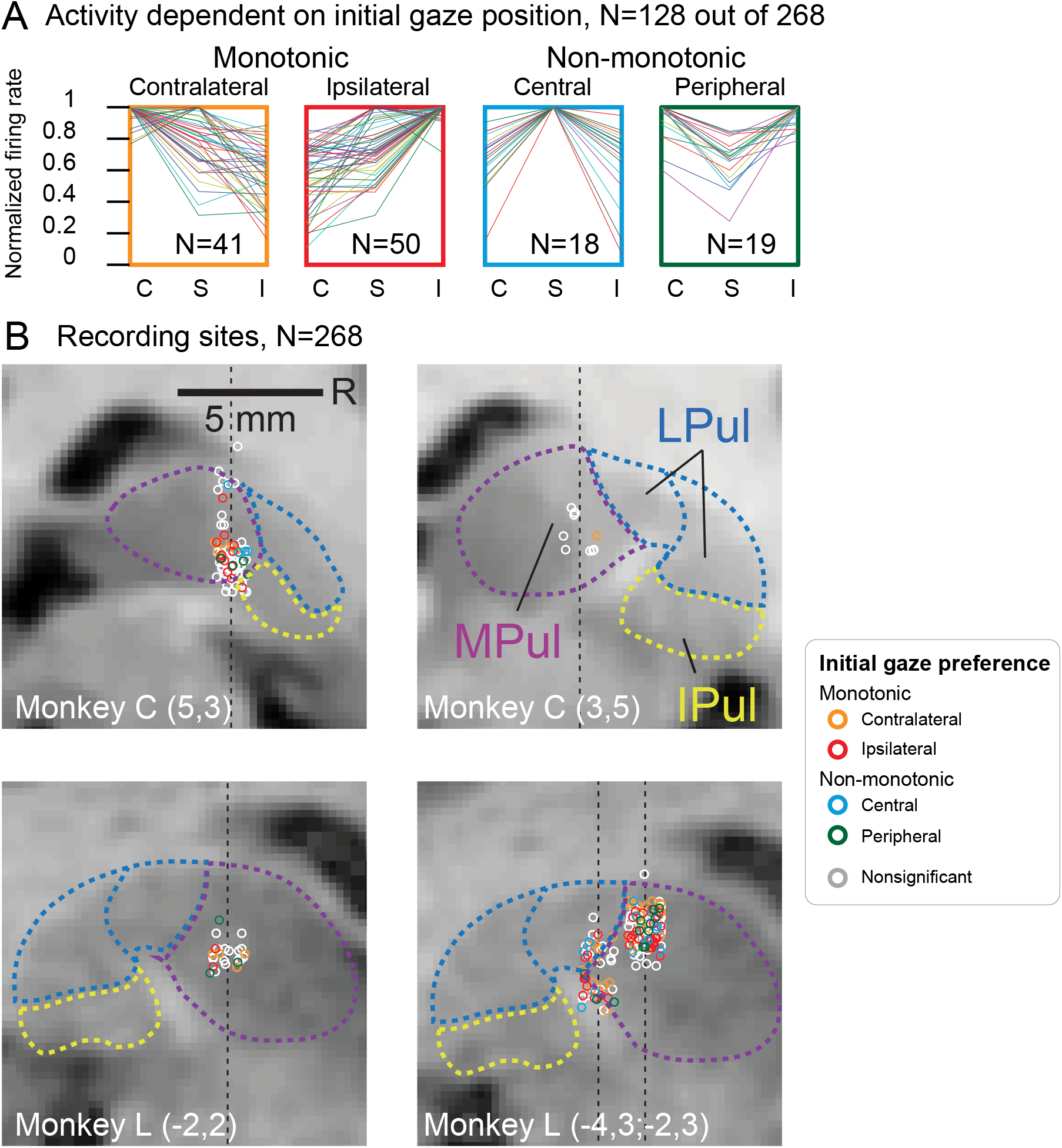
Gaze dependence during initial fixation and recording sites. **A:** Classification of initial gaze dependence. Normalized firing rates of each unit with a significant main effect of gaze on firing rates in the initial fixation epoch. Each line represents one unit. **B:** Localization of recorded units in chamber-normal coronal sections in each monkey (L and C, labels on the bottom) and specific grid locations relative to the chamber center (x,y in parentheses). Locations were jittered along the horizontal dimension for better visualization. The black dashed lines indicate the projection of penetration tracks and mark the actual horizontal location of recorded neurons. Each dot represents one unit; colors indicate the initial gaze effects of the unit: orange for monotonic contralateral preference, red for monotonic ipsilateral preference, blue for central gaze preference, and green for non-monotonic peripheral preference. Units that did not show a significant effect of gaze are in white. Pulvinar nuclei outlines (MPul/LPul/IPul – medial/lateral/inferior pulvinar) were adapted from the NeuroMaps atlas (Rohlfing et al., 2012), exported via Scalable Brain Atlas, https://scalablebrainatlas.incf.org/macaque/DB09, https://scalablebrainatlas.incf.org/services/rgbslice.php, (Bakker et al., 2015), and LPul was further subdivided into dorsal (PLdm) and ventral (PLvl) parts according to the brachium of the superior colliculus.

### Relationship between initial and final gaze effects

Next we evaluated the relationship between gaze-dependent activity during initial fixation and final fixation after the saccade (during target hold). Referring back to the examples in Figure 2, the top example shows mostly consistent contralateral gaze preference throughout the trial. The middle example shows consistent ipsilateral gaze preference. The bottom example shows inconsistent gaze preference (contralateral for the initial gaze and ipsilateral after the saccade). To evaluate these patterns across the population, we compared gaze position preference in fixation hold and target hold epochs (Figure 4A). Gaze position preference here was computed as the difference of firing rate averages for contralateral and ipsilateral gaze positions, see schematics in Figure 4A and Materials and Methods. Across all units gaze position preference in the two epochs was positively correlated (R=0.3, p<0.001, Pearson’s correlation), indicating a consistent effect of gaze direction irrespective of the task epoch. As expected, the correlation was mainly driven by units that show both an effect of initial and final gaze (significance derived from two independent one-way ANOVAs; N=91, R=0.4, p<0.001). Figure 4B shows the results of the two independent ANOVAs. Only 29 units showed an effect of initial gaze but not final gaze position; conversely, 65 units showed only an effect of final gaze position. The high number of units with only an effect of final gaze might be due to a wider range of (final) gaze positions in the target hold epoch.

**Figure 4.**
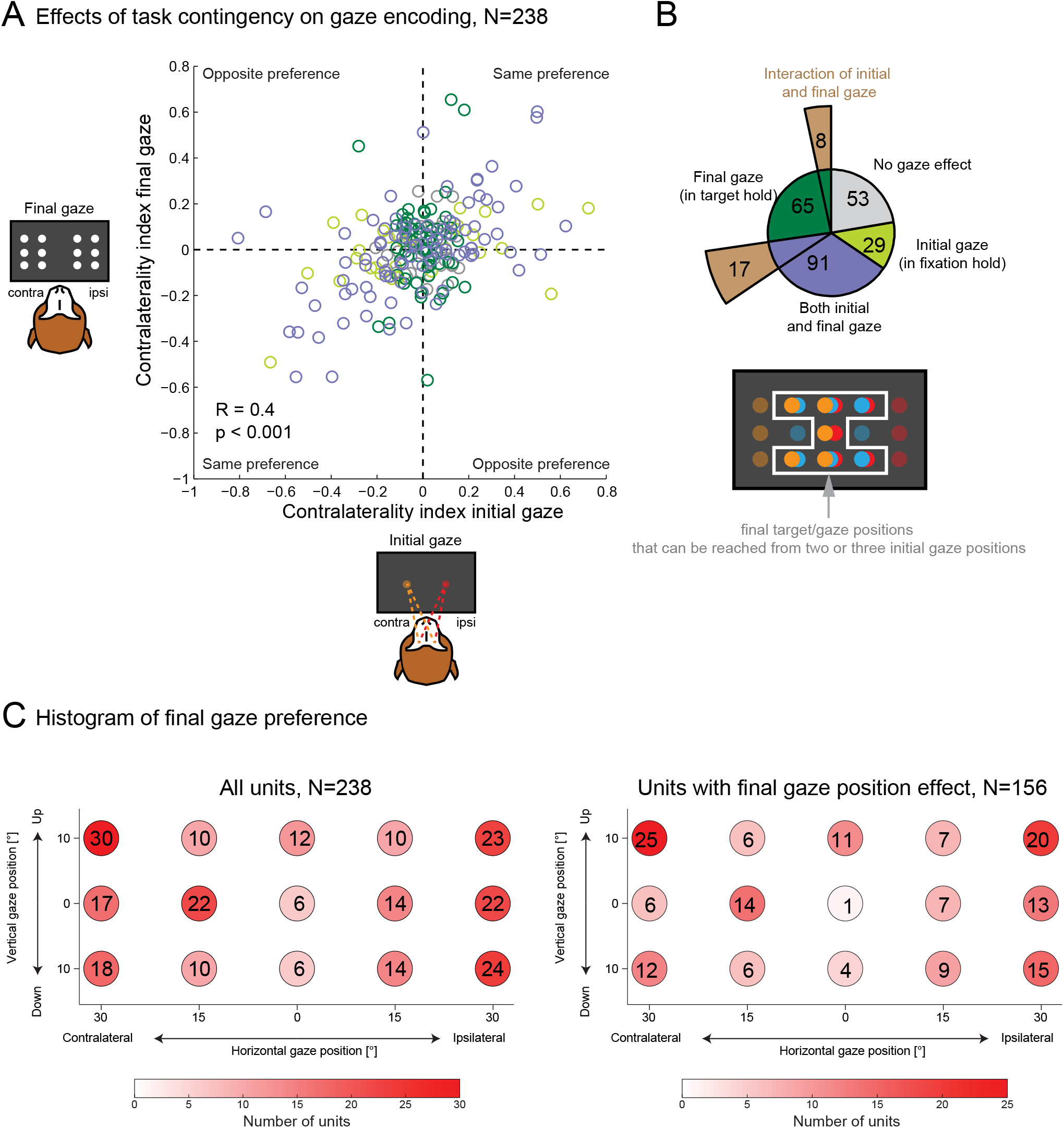
Initial vs. final gaze-dependent activity. **A:** Scatter plot of contralaterality indices denoting gaze direction preference in initial fixation (Fhol) vs. final target hold (Thol) epochs. Each circle represents a unit with at least 4 trials for every combination of initial gaze and retinocentric target position (N=238). Contralaterality indices were computed as difference of firing rates for contralateral and ipsilateral gaze positions, normalized by their sum. For the final gaze positions, 6 gaze positions in each hemifield were averaged, see schematic on the left. Color of each unit depicts the combination of effects: significant initial gaze effect only (lime), significant final gaze only (dark green), both initial and final gaze (purple), or none (gray). Effects of initial and final gaze position were defined by two independent ANOVAs, testing firing rates in Fhol for initial gaze position and firing rates in Thol for final gaze position. **B:** ANOVA results. Inner sectors: number of units in each group (same colors as in A), derived from the two independent one-way ANOVAs (on initial and final gaze). Outer sectors: number of units showing a main effect of final gaze as well as an interaction effect of the initial and the final gaze in the additional two-way ANOVA that used only the highlighted (white border) target positions depicted in the schematic below (brown). **C:** 2D histogram of final gaze preferences. Red color intensity indicates the count of units which showed the strongest response for the respective final gaze position. Left panel: all units, right panel: only units showing significant effect of final gaze position.

Next, we asked whether there is any influence of the initial gaze on activity associated with the final gaze after saccade. We performed a two-way ANOVA with the factors initial gaze and final gaze on firing rates in the target hold epoch, using the seven final target positions where the gaze would arrive from multiple starting points (schematic in Figure 4B). The outer sectors of Figure 4B display the number of units that showed a main effect of the final gaze and an interaction between initial and final gaze. Only 16% of units that showed an effect of final gaze position in the one-way ANOVA also showed interaction of initial and final gaze in the two-way ANOVA (25/156), indicating that trial history (e.g. preceding saccade direction) did not have a major impact.

Since the final gaze covered more spatial locations, we assessed final gaze-dependent firing rate preferences in 2D space. Figure 4C shows color-coded histograms of gaze positions associated with maximum firing rates. Substantially more units preferred peripheral horizontal gaze positions as compared to central gaze positions (chi-square test, p=0.00021 for all units, p= 0.00235 for significant units). This pattern is in agreement with the overall peripheral preferences during the initial fixation (cf. Figure 3A).

Given the sufficiently broad range of final gaze positions, especially in the horizontal dimension, we further quantified the monotonicity of the final gaze effects, separately for horizontal and vertical axes. We performed a two-way ANOVA with factors horizontal and vertical final gaze position on target hold firing rates, in all units that showed an effect of final gaze position in the one-way ANOVA (N=156). Table 1 shows the number of units that exhibited main effects of horizontal/vertical gaze position, further separated into monotonic and non-monotonic gaze preference. Most units (117/156, 75%) showed a main effect of horizontal gaze position, but only a minority showed monotonic gaze preference (32/117, 27%). Thus, the wider range of final gaze positions (as compared to only 3 initial gaze positions) revealed predominantly non-monotonic gaze dependence. Many units (87/156, 56%) showed a main effect of vertical gaze position, around two thirds of them showed monotonic vertical gaze preference (62/87, 71%). This indicates that the vertical gaze position had a strong impact on firing rates, but likely due to the smaller range of the vertical (compared to the horizontal) component less significant effects and more monotonic preference was detected. A more detailed picture of final gaze preferences for each unit can be gained from **Supplemental Figure S2**.

**Table 1.** Monotonicity in final gaze dependence. Number of units exhibiting an effect of final gaze position, grouped by significant monotonic and non-monotonic main effects of horizontal and vertical gaze position.

| Units showing an effect of final gaze position (N=156) |  | Horizontal axis |  |  |
| --- | --- | --- | --- | --- |
|  |  | Monotonic | Non-monotonic | Nonsignificant |
| Vertical axis | Monotonic | 13 | 26 | 23 |
|  | Non-monotonic | 5 | 19 | 1 |
|  | Nonsignificant | 14 | 40 | 15 |

### Gaze-dependent modulation of spatially-contingent task epochs

So far we addressed the effect of gaze position on the neuronal firing during either initial or final (target hold) fixation. Here we ask how the retinocentric spatially-contingent encoding is affected by gaze in the visual cue, memory delay and saccadic epochs. We performed a two-way ANOVA with factors initial gaze position and retinocentric target location for each unit and each epoch of interest, (Figure 5A). During cue presentation purely retinocentric encoding (only main effect of retinocentric target location) was less common then some dependence on the gaze position (41/238 units vs. 82/238 units), indicating a strong influence of current gaze position on the response strength or tuning pattern. In the memory and peri-saccadic epoch, purely retinocentric encoding became even less common (22/238 vs. 86/238 units and 25/238 vs. 50/238 units respectively). Finally, one third to half of the units showed an interaction of initial gaze position and retinocentric target location in the post-saccadic epochs (81/238 in the post-saccadic epoch and 114/238 in target hold epoch). For an approximate estimation of the false discovery rate due to random firing fluctuations, we performed the same analysis on firing rates in the initial fixation epoch (where no information about target location was available to the monkeys). While nearly half of the units showed only gaze-dependent activity during the initial fixation (108 out of 238), only a small fraction showed a main effect of target location only (10/238 units), both main effects (6/238 units), or any combination with [initial gaze × target position] interaction (11/238 units), as expected.

**Figure 5.**
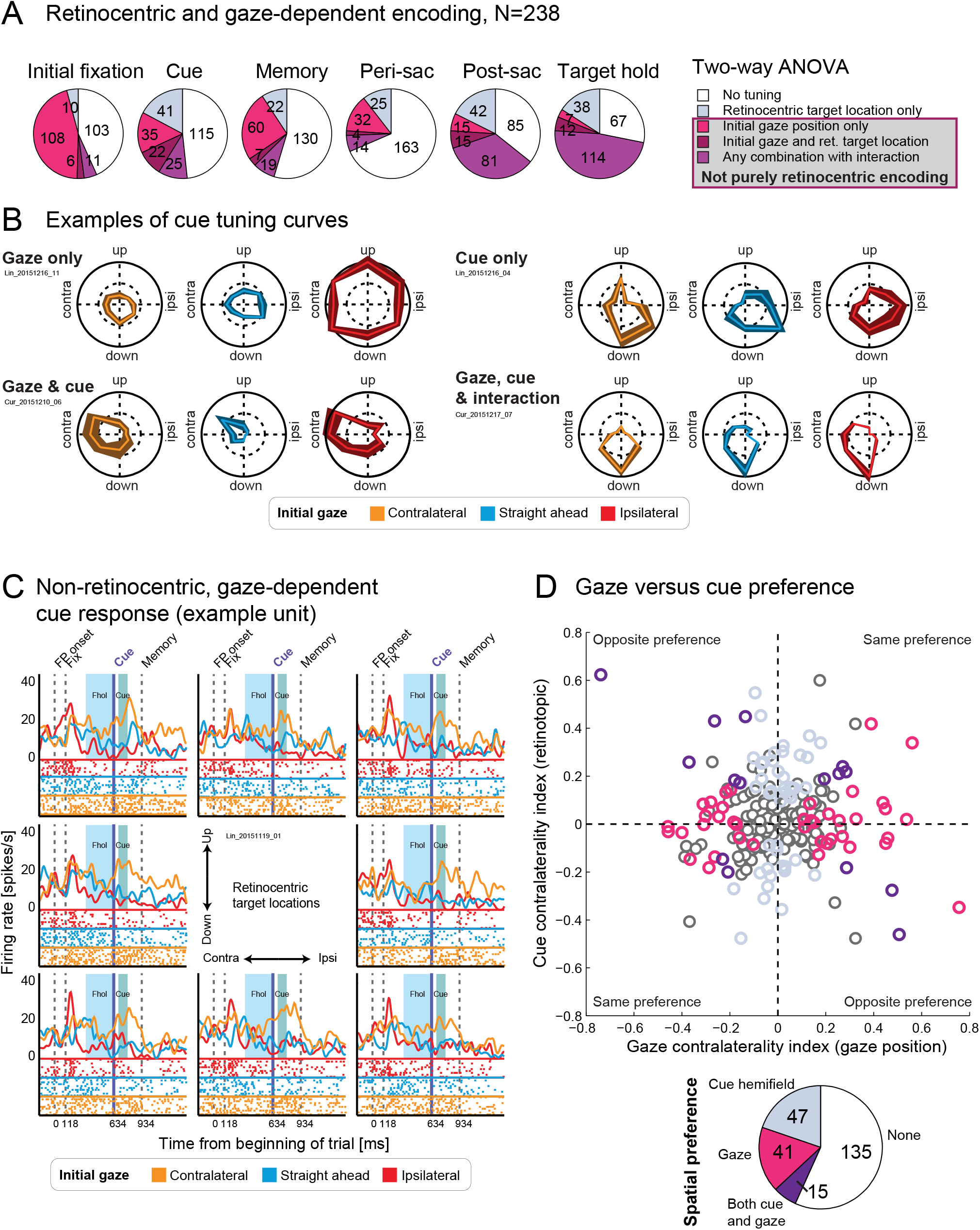
Retinocentric and gaze-dependent encoding. **A:** Two-way ANOVA results, with factors initial gaze position and retinocentric target location, independently for six epochs: fixation hold (Fhol), cue onset (Cue), memory (Mem), peri-saccadic (Peri), post-saccadic (Post), and target hold (Thol). Number of units showing either only a main effect of retinocentric target location, only a main effect of initial gaze position, both main effects, or any combination with [initial gaze position × retinocentric target location] interaction (plus either no main effect, both main effects, or one of the two main effects). For brevity, pre-saccadic epoch is omitted because the numbers were nearly identical to peri-saccadic epoch, target onset epoch is omitted because the numbers were nearly identical to post-saccadic epoch. **B:** Polar plots of retinocentric cue responses for each gaze position. One example unit from each category in the two-way ANOVA. **C:** PSTHs and raster plots of activity around the cue onset for one example unit showing a main effect of gaze (in the cue epoch) only, separately for each retinocentric cue location (in different subplots) and each gaze position (in different colors). **D:** Top: Scatter plot of cue and initial gaze contralaterality indices. Colors indicate significant hemifield preferences (light blue: cue, magenta: gaze, purple: both cue and gaze, grey: no hemifield preferences). Bottom: pie plot showing the number of units with hemifield preferences for cue, gaze, both or none.

The fraction of units with gaze dependence in the cue epoch decreased as compared to initial fixation (120 vs. 82 units). Therefore we asked if the arrival of the new spatial information overrode the initial gaze encoding in some neurons. Out of 120 units that showed an effect of gaze position during initial fixation, 53 units maintained gaze dependence (main effect of gaze or interaction) in the cue epoch (and additional 29 units acquired gaze dependence). Out of 67 units that did not maintain gaze dependence, only 23 showed a main effect of retinocentric cue position, meaning that for the other 44 units the loss of gaze encoding cannot be explained by replacement with new spatial cue information. Conversely, 22 out of 53 units maintained gaze dependence despite additional encoding of the new retinocentric cue position. This together indicates that the decrease in gaze dependence was not related to cue responses. We also asked, more generally, if the units with the initial gaze effect are less likely to have visual cue responses. Table 2 shows that it is not the case – units with and without an effect of initial gaze position exhibited similar cue response patterns.

**Table 2.** Relationship between initial gaze effect and cue responses. Number of units showing enhancement, suppression, hemifield preference, or an effect of retinocentric position during cue presentation, and none of the above, separately for units showing an initial gaze effect (in fixation hold, 120 units, cf. Figure 4B, 120=91+29) and units showing no effect of initial gaze (118). Please see the analysis of the relationship between initial gaze effect and saccade response in **Supplemental Table S1**.

| Fixation hold |  | Cue response |  |  |  | No cue response |
| --- | --- | --- | --- | --- | --- | --- |
|  |  | Enhancement | Suppression | Hemifield preference | ANOVA position effect |  |
| Initial gaze effect present | 120 | 28 | 25 | 33 (25 with ANOVA position effect) | 45 (19/10 en/su) | 50 |
| Initial gaze effect absent | 118 | 23 | 22 | 29 (17 with ANOVA position effect) | 30 (13/9 en/su) | 57 |

To get a better understanding of how gaze position affected visual responses, we looked into the cue epoch more closely. Example tuning curves for each category illustrate the combinations of gaze and cue dependence defined by the ANOVA (Figure 5B). Some units showed a strong modulation by gaze position but no directional cue tuning (“Gaze only”), some showed the similar retinocentric cue tuning regardless of gaze position (“Cue only”), some showed directional cue tuning scaled by gaze (“Gaze and Cue”), and some showed an alteration of preferred direction by gaze (“Gaze, Cue, and interaction”).

It should be noted here that some units showed a clear response to the cue onset specifically for one gaze position, but no spatial cue tuning, as illustrated in the example shown in Figure 5C. This finding demonstrates that cue responses may be underestimated when looking at only one gaze position, and should be considered in further studies assessing functional significance of cue responses in dorsal pulvinar.

Next we tested if there is a relationship between gaze and spatial cue preference in units showing both types of dependence. We found no correlation between cue and gaze spatial preferences, neither when correlating cue contralaterality indices (CIs) to gaze CIs derived from cue epoch (R=-0.04, p=0.513, Pearson’s correlation; see Figure 5D), neither when correlating cue CIs with gaze CIs derived from the fixation hold epoch (R=-0.05, p=0.471, Pearson’s correlation), nor when correlating raw (contra-ipsi) firing rate differences (R=0.05, p=0.478 for gaze in cue epoch; R=-0.03, p=0.619 for gaze in the fixation hold epoch). In fact, many units showed either hemifield preference of gaze position (N=56) or retinocentric cue location (N=62), but only few units showed both (N=15), with only 6 units showing the same hemifield preference.

To compare the strength of cue and gaze tuning, we computed absolute gaze and cue CIs and raw (contra-ipsi) firing rate differences and tested if there was a difference of the mean ranks (Wilcoxon signed rank test). There was no significant difference between the strength of cue and gaze tuning in any of the comparisons mentioned above (p=0.103 for cue CIs and gaze CIs in cue, p=0.104 for cue CIs and gaze CIs in fixation hold, p=0.2 for raw firing rate differences (gaze in cue), p=0.295 for raw firing rate differences (gaze in fixation hold).

### Modulation of retinocentric tuning by gaze position

In this section we focus on how gaze direction modulated units that showed retinocentric tuning. The motivation is to assess whether gaze-dependent modulation would be consistent with having gaze-dependent gain fields. To this end, we bootstrapped the tuning vector (dependent on retinocentric target location) for each initial gaze position, independently for each unit and each epoch. To assess if gaze position amplified or shifted response fields, we computed confidence intervals for the estimated tuning vector for each gaze position and compared the confidence intervals around the mean for vector length and direction, see Materials and Methods. This approach allowed us to assess changes in tuning without reliance on a specific tuning function. To illustrate this procedure, we modelled a neuron with a Gaussian retinocentric response field and a gaze-dependent linear multiplicative gain field and simulated recorded data for this hypothetical neuron by adding random noise for each trial (see Figure 6A). Using this simulated data for bootstrapping the tuning vector for each gaze position and comparing confidence intervals of estimated tuning vector end points confirmed that the gain field properties of the modelled neuron could be reconstructed (see Figure 6B).

**Figure 6.**
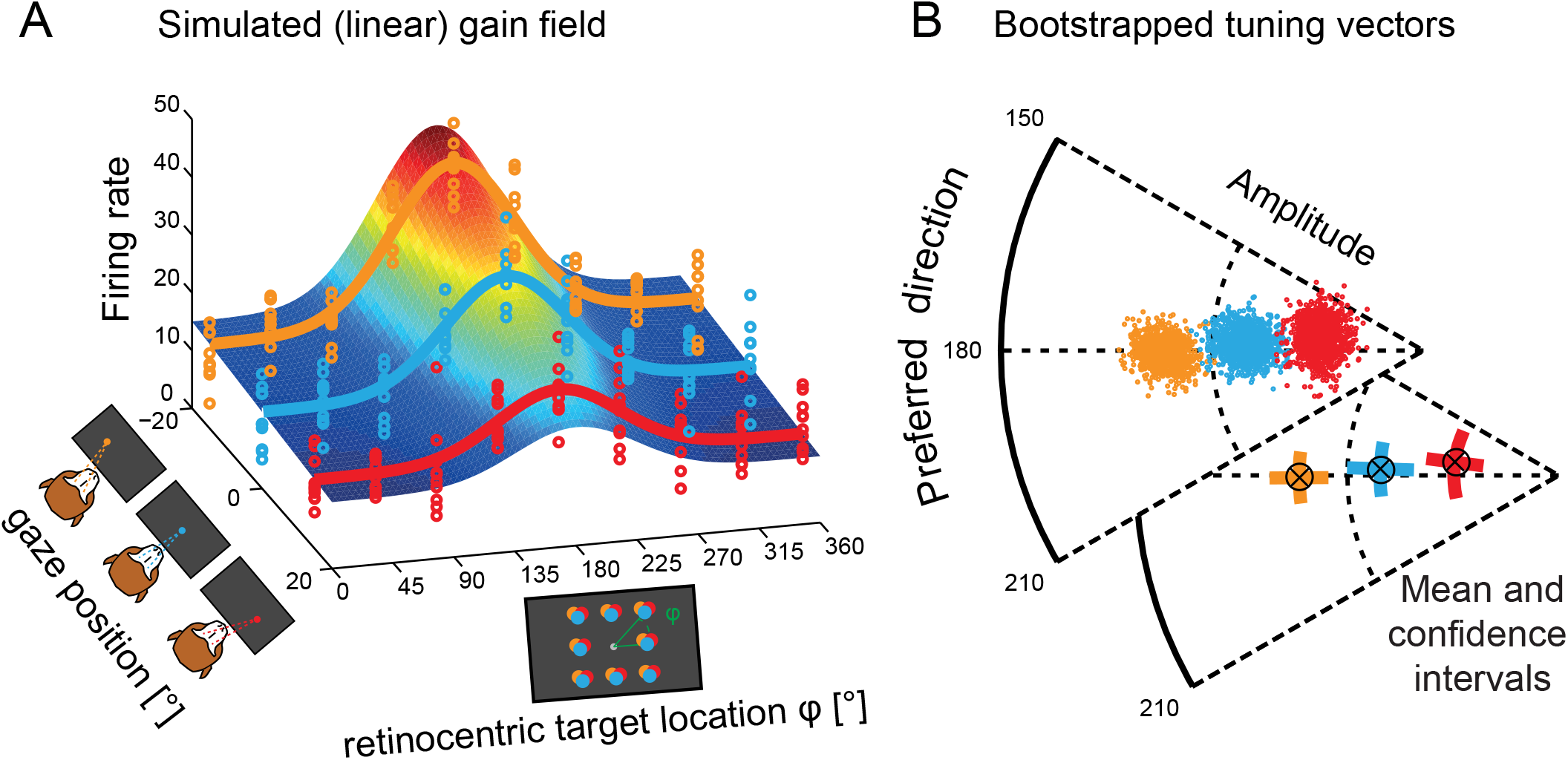
Tuning vector bootstrapping approach. **A:** 3D plot of a simulated gain field. Each color corresponds to one initial gaze position. Firing rates were simulated using a Gaussian tuning curve (dependent on retinocentric target location, preferred direction = 180°) which was amplified by the initial gaze position. Noise was added to create variability for each trial (colored circles). To illustrate the comparison with the results derived from the two-way ANOVA with factors retinocentric location and gaze position, such gain field results in both main effects and interaction. **B:** Population vectors for each initial gaze position were bootstrapped (see Materials and Methods) to compute confidence intervals for amplitude and direction of the estimated population vectors for each initial gaze position.

Spatial firing rate patterns of six recorded example units exhibiting either gain field properties or response field shifts in the cue epoch are shown in Figure 7A. For further illustration, PSTHs for each combination of initial gaze and retinocentric target location as well as the distribution of bootstrapped tuning vector endpoints are shown for the two top example units. The left example shows a classical gain field, retinocentric cue response amplified by gaze position. The example on the right shows a more complex pattern: here the preferred direction is modulated by the gaze position, with more ipsilateral gaze amplifying the central and upper contralateral cue response.

**Figure 7.**
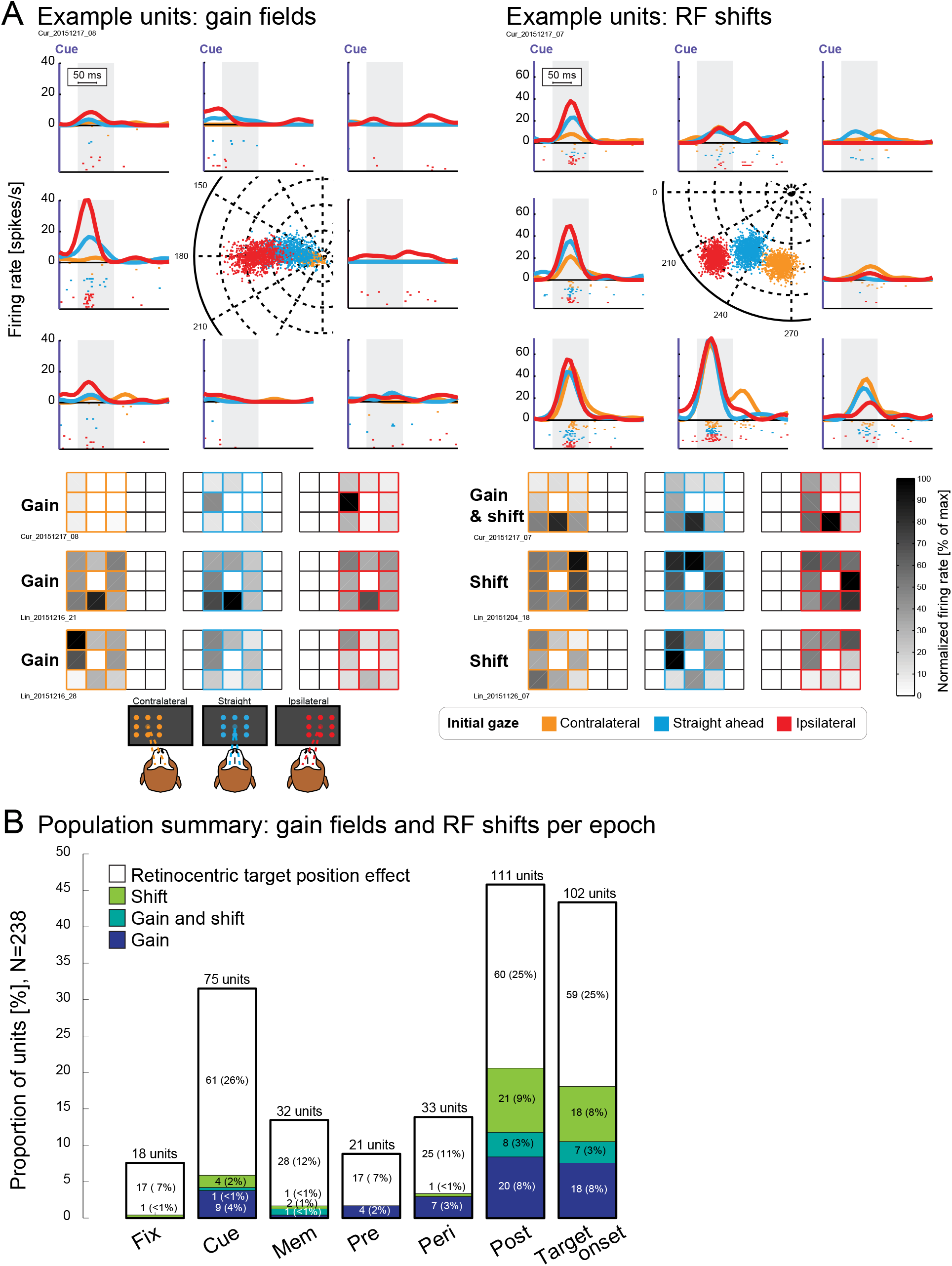
Gain field and response field (RF) shift properties. **A:** Example units showing gain field properties (left) or RF shifts (right) in the cue epoch. Top: PSTHs during cue presentation for each of the initial gaze positions and each of the retinocentric target locations, spatially arranged according to the retinocentric location. In the center of the PSTH plots, the corresponding bootstrapped population vector endpoints for each initial gaze position are displayed. Bottom: Heat map plots of the example units shown above (top) and two additional examples showing either gain field properties or RF shift properties. **B:** Number of units showing a main effect of retinocentric target position (top of the bar), and out of those, units showing gain field (blue) or RF shift properties (green) for each epoch of interest. Units showing both effects are displayed in light blue, units showing only an effect of target position are displayed in white. Fixation hold (Fix) epoch was added as a control in order to approximate the expected statistical noise level.

Figure 7B shows the number of units exhibiting a main effect of retinocentric target location (derived from two-way ANOVA in Figure 5A), and out of those units, the number of unit showing gain, shift or both, for each epochs. Among the epochs occurring prior to the saccadic eye movement, and thus reflecting the potential effects of the static gaze, we focus on visual cue responses. Out of 75 units that showed the main effect of cue/target location, only 10 units showed a significant gain field component (7 monotonic). In those units, the gain amplitude was 66±15% (mean±SD, Materials and Methods). Importantly, our gain field analysis only detects multiplicative impact of gaze on retinocentric tuning, which in the two-way ANOVA analysis (cf. Figure 5A) would be reflected by a combination of the main effect of retinocentric target location and interaction. Indeed, 9 out of the 12 units showing this combination were identified as showing significant gain field. The 22 units which showed main effects of retinocentric location and gaze but no interaction might be attributed to additive effects which are not defined as gain. The additive effect could be illustrated by the tuning curves of the example unit in Figure 5B (Cue and Gaze).

The interpretation of the two post-saccadic epochs (immediately after saccade, and during target onset) is more complicated because gaze-dependent effects could also reflect the final gaze position (which is not fully dissociated from the initial gaze position). In fact, 46 out of 53 units showing gain and/or shift in the post-saccadic epoch and 42 out of 45 units showing gain and/or shift in the target onset epoch also show an effect of final gaze position in the same epoch, making gaze position encoding the most parsimonious explanation for the relatively frequent gain and shift effects in these epochs.

As a control, we also performed this analysis on firing rates during fixation hold to get an estimate for the false discovery rate due to random fluctuations (since at that point there was no difference between trials with different retinocentric target locations). Only one unit showed a spurious “shift” effect, suggesting that the findings in other epochs are not due to random noise fluctuations.

### Reference frame evaluation

Finally, we evaluated in which reference frame dPul neurons encode spatial information. Since the monkey’s head position was always immobilized straight ahead relative to the screen, and we did not control for the trunk rotation in the chair (although typically the trunk was facing the screen), we cannot dissociate between head-, body-, or world-centered representations. However, we can dissociate these three possibilities from retinocentric (i.e. eye-centered) representations. For simplicity, in the following text a “screen-centered” reference frame refers to a potential head-, body-, or world-centered reference frame, as opposed to a retinocentric one.

To evaluate if response patterns were better explained by retinocentric or screen-centered representations, we compared the alignment of responses in retinocentric and screen-centered coordinates for seven task epochs, following a previously applied approach (Materials and Methods, (Mullette-Gillman et al., 2005)). For this analysis, we included only units that exhibited a dependence on target locations in any reference frame: i.e. only units which showed either a main effect of retinocentric target location, [retinocentric location × initial gaze] interaction, a main effect of target location on the screen, or [screen location × initial gaze] interaction. In Figure 8, each data point represents the average correlation coefficient (ACC) between the unit’s responses for different initial gaze positions when locations were defined relative to the eye (retinocentric ACC, horizontal axis) and relative to the screen (screen-centered ACC, vertical axis). Units above the diagonal indicate better alignment of responses in screen-centered coordinates, and units below the diagonal indicate better alignment in retinocentric coordinates. Units showing response patterns that are significantly better explained by a screen-centered reference frame are in green, units that are better explained by a retinocentric reference frame are in red. Figure 8 shows a progression from prevalence of units compatible with retinocentric reference frames in the cue epoch to screen-centered encoding in the target hold epoch, with a balanced representation immediately after the saccade. This indicates that visual responses and/or movement preparation were encoded in retinocentric coordinates, while the new gaze position was the predominant factor encoded in post-saccadic epochs. As additional support for this interpretation, we performed paired t-tests comparing retinocentric ACCs and screen-centered ACCs across all included units in each epoch, showing significantly stronger retinocentric alignment in the cue (p<0.001), memory (p=0.01) and peri-saccadic (p=0.001) epochs and stronger screen-centered alignment in the target onset (p=0.003) and target hold (p<0.001) epochs.

**Figure 8.**
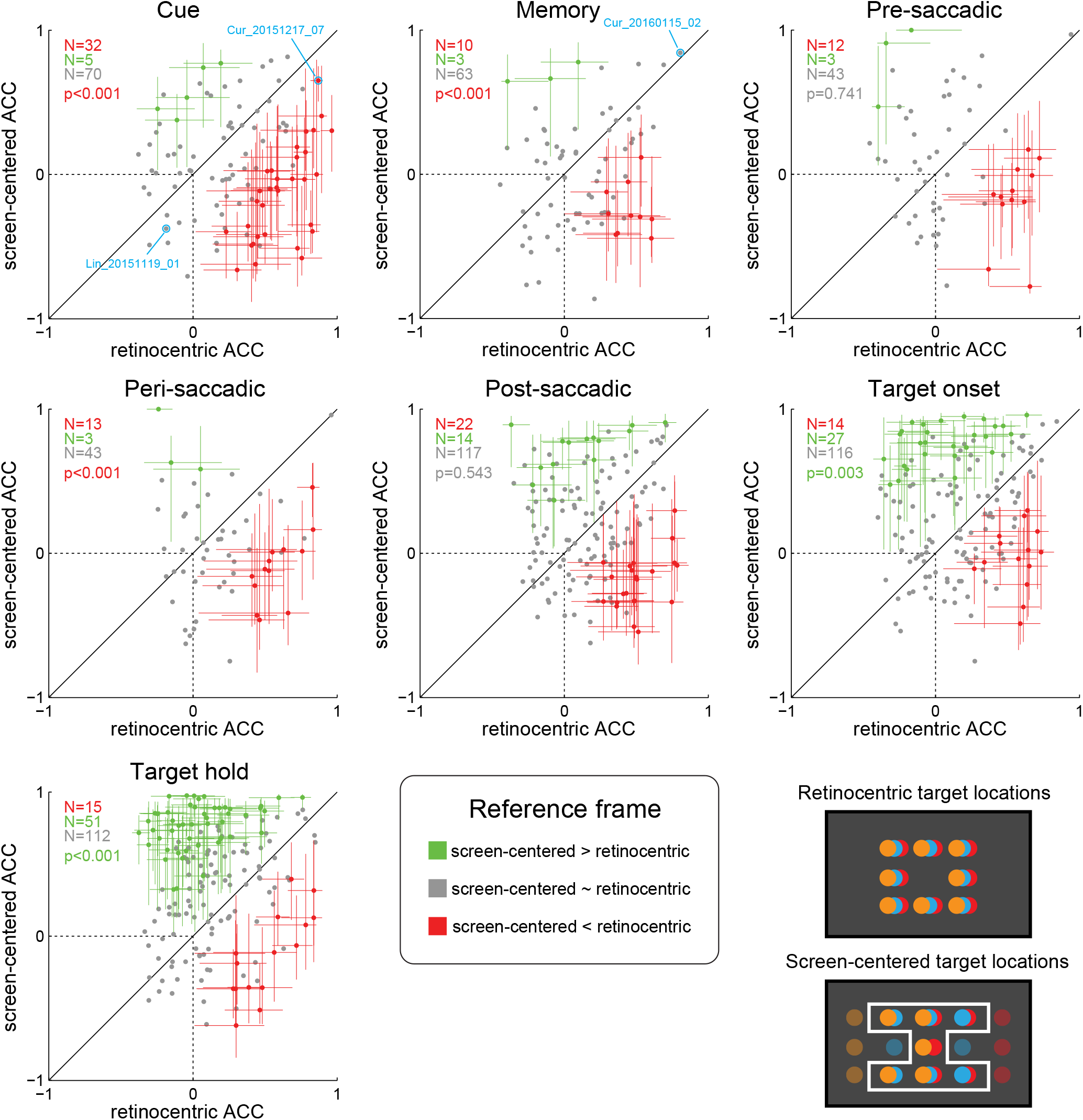
Reference frame evaluation. Average correlation coefficients (ACCs) in retinocentric coordinates (x-axis) and screen-centered coordinates (y-axis) for each unit in seven epochs of interest: cue, memory, pre-saccadic, peri-saccadic, post-saccadic, target onset and target hold epoch. The crosshairs indicate 95% confidence intervals derived from bootstrapping. Note that dots represent actual ACCs calculated using all data, and their location is not necessarily centered in the middle of the confidence intervals. Units where the crosshair is above the diagonal (and >0 on the y-axis) were classified as “more screen-centered than retinocentric” (green) and units where the crosshair is below the diagonal (and >0 in the x-axis) were classified as “more retinocentric than screen-centered” (red), see Materials and Methods for details. For the remaining units (mixed, or hybrid encoding, gray), confidence intervals were omitted for clarity. For eye-centered correlations, all 8 target locations were used, for screen-centered correlations, only overlapping target locations for each specific correlation pair were used (7 targets outlined by the white frame). P-values indicate significance of paired t-tests comparing screen-centered and retinocentric ACCs across all included units, the text color indicates which one was larger (in case of significant differences). For cross-referencing, we marked two units in the cue epoch; the example from Figure 5C (with main effect of gaze but no effect of retinocentric location, Lin_20151119_01) showing negative correlation in both reference frames, and the example from Figure 7A right (showing gain and shift, Cur_20151217_07), which shows a strong correlation in both reference frames (but significantly more for retinocentric), as well as one unit for the memory epoch, the example from **Supplemental Figure S3** (Cur_20160115_02), a mixed unit which shows a strong correlation in both reference frames, but no significant difference between them.

Apart from the units that could be classified as retinocentric or screen-centered, in each epoch, many units showed mixed, or hybrid encoding that cannot be more strongly attributed to either of the two references frames (Caruso et al., 2018). Many of these units had high ACCs in both frames, i.e. the both reference frames could account for the observed response functions. To illustrate this, we show an example unit that responded predominantly to upper targets, and had high ACCs in both reference frames (e.g. 0.84 for screen-centered and for 0.81 retinocentric in the memory epoch), making it difficult to dissociate the two (**Supplemental Figure S3**). This and other mixed units showed response patterns that are consistent with a partial shift of response function with the eye position (Caruso et al., 2018).

## Discussion

The dorsal pulvinar has been implicated in spatial processing supporting goal-directed eye movements, but so far the implicit experimental assumption was that it encodes space in retinocentric (eye-centered) coordinates, even if it does not show a retinotopic organization (i.e. gradual topographic representation of visual space across adjacent regions of neural tissue). Here we show that many dorsal pulvinar neurons are influenced by the position of the eyes in the orbit (equivalent to the gaze direction, when the animal’s head is immobilized), in two different ways. First, in more than half of the sample the firing during steady fixation of a foveal stimulus was influenced by the gaze direction. Second, depending on the epoch substantial numbers of neurons showed a combination of retinocentric and gaze-dependent signals in visual and saccadic responses to a peripheral stimulus, and some of these effects could be described as gaze-depended gain modulation. The analysis of reference frames across consequent task epochs, from cue to post-saccadic target hold, indicated a transition from predominantly retinocentric encoding to coding of final gaze or target location in head/body/world-centered coordinates. Despite some limitations in the current study design (see below), these results provide electrophysiological evidence to support the previously hypothesized notion that the dorsal pulvinar might contain neurons that reflect postural signals and represent space in non-retinocentric reference frames (Grieve et al., 2000). In the following sections, we relate our findings to the previous cortical and thalamic literature, separately discussing the influence of eye position on ongoing firing during foveal fixation, and on visuomotor representations of peripheral targets.

### Comparison to previous electrophysiological studies in the thalamus

Previous electrophysiological studies in the central thalamus of monkeys have reported neural activity modulation by eye position in the orbit (Schlag-Rey and Schlag, 1984; Wyder et al., 2003; Tanaka, 2007). Similar to the dorsal pulvinar, the central thalamus is a ‘higher-order’ nucleus complex with connections to cortical regions such as frontal and supplementary eye fields and posterior parietal cortex, that are crucial for oculomotor behaviors (Tanaka and Kunimatsu, 2011). The central thalamus also receives afferent inputs from the cerebellum and oculomotor brainstem involved in eye movement control.

More specifically, in respect to effects of eye position on ongoing fixation activity, a study in the central (‘oculomotor’) thalamus that investigated a number of nuclei such as paracentral nucleus (PC), centrolateral nucleus (CL), ventral anterior nucleus (VA), ventral lateral nucleus (VL), a paralaminar centromedian nucleus (CM) and paralaminar mediodorsal nucleus (MD), reported a small subgroup of 6 neurons showing an influence of gaze on sustained post-saccadic activity, either increasing or decreasing monotonically with the contralateral gaze eccentricity (Wyder et al., 2003). Comparing these results to the effects of the final (post-saccadic) gaze position in our sample, the majority of dPul units did not show monotonic dependence (e.g. for the 5 gaze positions on the horizontal axis, 32 units showed a monotonic effect and 85 were non-monotonic). The number of neurons showing monotonic gaze effects was much higher (71%) for the initial fixation, but this estimate is limited by only three gaze positions covering a relative narrow range (30°).

Another study that specifically focused on eye position responses in the central thalamus also found, in addition to transient post-saccadic responses, many neurons with sustained post-saccadic firing modulated by eye position (Tanaka, 2007). Most neurons in this study had positional preferences along the horizontal axis. Some neurons also showed a “memory trace” of the preceding saccade, i.e. the activity at the same final position depended on where the eye came from. Such hysteresis effect was also reported in the central thalamus during spontaneous eye movements (Schlag-Rey and Schlag, 1984). This is reminiscent of our subset of neurons showing an interaction between the initial and the final gaze position. But unlike our findings, the eye position effects in the central thalamus could be divided into the two main groups: the neurons that showed predictive pre-saccadic tuning corresponding to the final gaze position, and those that were modulated by the gaze only after the saccade (Tanaka, 2007). In our sample, the majority of neurons were similar to the latter, while the units showing pre-saccadic responses consistent with encoding the post-saccadic gaze were exceedingly rare.

The thalamus literature on gaze-dependent modulation of peripheral (eccentric) visual responses (e.g. potential gain field effects) is sparse. The modulation of visual responses by the (passive) eye position has been reported in lateral geniculate nucleus (LGN) of anesthetized cats (Lal and Friedlander, 1989). The eccentric visual responses in the cat internal medullary lamina (IML) were found to be related to a confluence of specific retinocentric direction and a target area defined in screen/head/body-related coordinates, determined by spontaneous gaze locations (Schlag et al., 1980). In the retinotopically organized macaque ventral pulvinar, visual responses of a subset of neurons were influenced by orbital position (Robinson et al., 1990). In this study, stimuli were briefly flashed in the receptive field at different times around visually-guided saccades, to assess the visual sensitivity dynamics. In 15 out of 41 cells, modulation of visual sensitivity by eye position has been detected. Apart from those pioneering studies, to our knowledge there has been no systematic assessment of potential gaze-dependent gain fields in higher-order thalamus.

### Comparison to previous electrophysiological studies in cortex

Eye position effects in cortex were mostly considered in the context of reference frame transformations and integration of sensory information into head- and body-centered coordinates. Nonetheless, the studies that have tested the influence of gaze on ongoing activity during fixation in task periods unrelated to the representation of peripheral targets for goal-directed movements (even in the dark) also reported eye position-dependent effects. Specifically, eye position signals during initial fixation have been reported in many cortical areas including visual (V1-V4, medial temporal area MT, MST) (Bremmer, 2000; Rosenbluth and Allman, 2002; Morris et al., 2013), posterior parietal (LIP, MIP, 7a, VIP, V6) (Sakata et al., 1980; Andersen and Mountcastle, 1983; Andersen et al., 1990; Galletti et al., 1995; Bremmer et al., 1997; Mullette-Gillman et al., 2005; Genovesio et al., 2007), dorsal premotor (Boussaoud et al., 1998) and somatosensory cortex (Wang et al., 2007). Below we briefly discuss a number of particularly pertinent studies.

Sakata et al. detailed eye position effects of neurons recorded from parietal area 7a that were clearly driven by gaze and did not show activity during passive visual stimulation (‘fixation neurons’). In their sample, ∼10% of neurons exhibited this property (Sakata et al., 1980). The majority of the 86 reported neurons varied systematically with gaze angle in the frontal plane (39 horizontal, 38 vertical, 9 diagonal). Many of those neurons were selective for depth of fixation as well, an aspect that is important but was not measured in our current study. The results of Sakata and colleagues are broadly consistent with later studies in areas LIP and 7a, where subpopulations of neurons were reported to exhibit tonic background activity during fixation that varied monotonically with eye position without showing any visual responsiveness (Andersen et al., 1990). Similarly, another study in LIP and 7a reported an influence of eye position during visual fixation in darkness with about a third of neurons that could be best fit with a monotonic (linear) regression model (Bremmer et al., 1997). A comprehensive study that compared how well eye position signals during fixation can be decoded from neurons recorded from LIP, VIP or visual areas MT and MST concluded that eccentric fixations could be decoded more accurately than central ones and that LIP/VIP populations encode eye position more accurately than visual cortex (Morris et al., 2013). Eye position signals were also reported in areas that are not primarily associated with visual or saccade processing. For example, a subpopulation of neurons in premotor area PMd were reported to differentiate between gaze angles in a reach task without being tuned to visual inputs or movement directions (Boussaoud et al., 1998).

Eye position signals that were monotonically related to the eccentricity of gaze were also reported in primary somatosensory cortex S1 (Wang et al., 2007; Xu et al., 2011). This study concluded that those eye position signals in S1 had a proprioceptive origin since anesthesia of the contralateral orbital muscles nearly abolished the recorded eye position signals. Similarly, a preference for more eccentric eye positions was reported in parietal cortex (Sakata et al., 1980). On the other hand, some studies in parietal cortex reported a fairly balanced distribution of gaze-depending activity on the population level (Bremmer et al., 1998). In the dPul we found a predominance of more eccentric (peripheral) gaze preferences, both during initial fixation and during target hold.

While the relationship between gaze-dependent effects during fixation and peripheral target encoding has been rarely explicitly discussed, it can be inferred from studies in parietal and premotor cortex that all three categories are typically observed: namely i) neurons that are modulated by the gaze during fixation but are not responsive in subsequent task epochs, ii) neurons that only show modulation of visual/delay/saccade responses by gaze, and iii) neurons that show both types of gaze-dependence. Similarly, we found all three categories in dPul, and no systematic relationship between presence or absence of gaze dependence and visual cue responses.

The majority of studies on eye position signals in cortex focused on the second aspect of gaze dependence: modulation of visual, motor planning and saccade-related activity in response to peripheral targets (Lehky et al., 2016). Specifically, when a neuron changes its response to a stimulus depending on eye position while the retinal location remains the same, this pattern has been described as eye position gain field. Computational models applied to the response modulation of single neurons in parietal (Andersen et al., 1990) and visual cortex (Bremmer, 2000) have often modeled the gain fields as planar, or at least monotonic (Zipser and Andersen, 1988; Pouget and Sejnowski, 1997). It needs to be noted however that even in posterior parietal cortex, the proportion of neurons reported to exhibit planar gain fields was only 39% (Andersen et al., 1985), and multidimensional scaling methods applied to neuronal populations in the lateral intraparietal cortex (LIP) and anterior inferotemporal cortex (AIT) to infer current eye position show that other, nonlinear functions (e.g. sigmoidal, elliptical or hyperbolic, or complex mixtures of thereof) might fit the data as well (Lehky et al., 2016). Likewise, eye position gain fields of neurons in area V6 (PO) have been shown to have diverse shapes from planar to peak-shaped (Galletti et al., 1995). Similarly to those findings, in the current study the gain fields in dPul appeared to be a mixture of monotonic and peak-shaped fields (but given only three initial gaze positions we used here, those conclusions are tentative). This is unlike the primary visual cortex, where the straight ahead direction typically stronger gain, leading to mainly peak-shaped fields (Durand et al., 2010; Przybyszewski et al., 2014).

In addition to the modulation of retinocentric encoding by eye position, several other oculomotor studies also reported encoding of visual/saccade targets in other reference frames, e.g. head-centered or mixed, or hybrid coordinates (Brotchie et al., 1995; Galletti et al., 1995; Mullette-Gillman et al., 2005; Caruso et al., 2018). Apart from a retinocentric reference frame, we also observed a small subset of neurons that were more consistent with head/body/world-centered encoding prior to the saccade, and many neurons with mixed encoding. For instance, some units showed (not tuned retinocentrically) visual cue responses only at certain gaze positions (cf. Figure 5C). Since the eye position could determine whether there was a visual response at all, it is conceivable to construe those neurons as primarily encoding eye position that is modulated by a visual response as opposed to the typical concept of eye position ‘gain fields’ where the visual response is modulated by eye position.

In our data, the prevalence of significant eye-centered encoding in cue, memory and pre-saccadic epochs is supplanted soon after the saccade by head/body/world-centered reference frame encoding, which can also be interpreted as the encoding of the current gaze. This finding is in line with ANOVA results showing the final gaze/target encoding during late post-saccadic fixation. The transition from retinocentric target to gaze position encoding in dPul resembles a recent study in FEF, with the difference that the transition happens after the saccade in dPul, as compared to the predictive pre-saccadic transition from retinocentric target code to (also retinocentric) motor goal in FEF (Sajad et al., 2016). The immediate post-saccadic epoch in dPul still contained units with retinocentric encoding, akin to eye movement vector post-saccadic encoding in LIP which dissipated in the late post-saccadic period (Genovesio et al., 2007). A small subset compatible with retinocentric encoding during target hold, long after the saccade, was unexpected, but the visual inspection confirmed that those units indeed retained a trace of the retinocentric saccade vector.

### Possible sources of eye position signals in dorsal pulvinar

Given the extensive connectivity of the dorsal pulvinar to parietal and premotor cortex, and the resemblance of eye position-dependent effects in those areas and in dPul, one likely possibility is that the eye position effects in dPul are inherited from those visuomotor areas (Yeterian and Pandya, 1989; Schmahmann and Pandya, 1990; Romanski et al., 1997; Cappe et al., 2007, 2009). The source of continuous eye position signal in the cortex is still not fully identified (Ziesche and Hamker, 2011; Xu et al., 2012). One possibility is the corollary discharge of the motor command that maintains steady eye position (Morris et al., 2012; Xu et al., 2012), counteracting the elastic elements of the orbital tissues (Sylvestre and Cullen, 1999). Another candidate is the proprioceptive signal reflecting the current position of the eyes in the orbit (Wang et al., 2007). The dPul can receive this signal from parietal and premotor areas, and/or directly from the primary somatosensory cortex (Wang et al., 2007; Xu et al., 2011), given medial pulvinar connectivity to area 3a (Padberg et al., 2009). This possibility could be investigated further by anaesthetizing the extraocular muscles (Wang et al., 2007), or by inactivating the projection from somatosensory area 3a and recording in the dPul.

Yet another possibility, which would place dPul as a potential source of eye position signals to the cortex, is the corollary discharge via superior colliculus – pulvinar – cortical pathway (Guthrie et al., 1983; Wurtz et al., 2011a). Dorsal pulvinar, including the lateral portion of the medial pulvinar where our recordings were mainly performed, receives afferents from the intermediate and deep layers of the superior colliculus (Harting et al., 1980; Benevento and Standage, 1983; Bender and Butter, 1987; Baldwin and Bourne, 2017). The intermediate and deep layers of SC exhibit eye position dependence, including gain fields (Van Opstal et al., 1995; Campos et al., 2006). It has been shown that besides the proprioceptive information about eye position (Andersen et al., 1990; Bremmer et al., 1997), a preparatory corollary discharge / efference copy of planned saccade, originating in the superior colliculus and routed to frontal cortex via mediodorsal thalamus (MD) can carry eye position signals (Crapse and Sommer, 2008; Wurtz et al., 2011a). Similarly, it has been suggested that the dorsal lateral pulvinar might carry preparatory movement-related signals to parietal cortex (Wurtz et al., 2011b), although those signals might only be relevant around the time of a saccade.

Although anatomical projections of the superior colliculus to the dPul have been reported (but see (Zhou et al., 2017)), there is a debate on specificity of projections to medial vs. lateral pulvinar and the functional impact of this projection remains unclear (Baldwin and Bourne, 2017)). The functional contribution of projections from SC is more studied in the ventral pulvinar, where the effects of SC perturbation or lesions have been assessed in several studies. Specifically, extensive lesions of SC caused only little effect on visual responses in the ventral pulvinar, as compared to striate cortex lesions (Bender, 1983). In rabbits, however, the inactivation of SC led to a strong attenuation of responses in lateral posterior nucleus (which is a part of the LP-pulvinar complex in non-primate species: mice, rats, rabbits, cats, and tree shrews) (Casanova and Molotchnikoff, 1990). Furthermore, selective microstimulation of the superficial layers of the SC in macaques has been shown to elicit (monosynaptic) responses in the ventral pulvinar (Kinoshita et al., 2019), in agreement with earlier work that identified the projection from SC to area MT via ventral pulvinar (Berman and Wurtz, 2011). Extrapolating from these ventral pulvinar studies to the dorsal pulvinar, the potential influence of SC inputs to dPul (Wurtz et al., 2005) might be outweighed by more extensive cortical driving inputs (Rovó et al., 2012; Bickford, 2016). To fully elucidate complex cortico-pulvinar-cortical loops and subcortical inputs to pulvinar, it would be crucial to manipulate the specific projections from and to the pulvinar, using pathway-selective techniques, such as optogenetics and/or viral vectors (Schmitt et al., 2017; Kinoshita et al., 2019).

Apart from cortex and superior colliculus, another possible source of eye position signals in the pulvinar is the cerebellum. The cerebellum is involved in a number of eye movement control functions including maintaining a steady fixation, smooth pursuit and binocular alignment (Patel and Zee, 2015; Krauzlis et al., 2017). To our knowledge no study in monkeys or humans has described (monosynaptic) pathways between dorsal pulvinar and cerebellum. In cats, several earlier histological labeling studies described projections from the cerebellum to the contralateral lateral-posterior nucleus (LP) of the thalamus, which is considered a putative homologue of the pulvinar (Itoh and Mizuno, 1979; Rodrigo-Angulo and Reinoso-Suarez, 1984). These cerebello-LP projections in cats were found to overlap with the ones coming from the deep layers of the superior colliculus and other brainstem nuclei. It is not clear however whether these results can be extrapolated to the medial portion of the dorsal pulvinar, which is only fully developed in primates (Jones, 2007; Preuss, 2007). Some evidence (albeit not necessarily for a monosynaptic pathway) can be derived from microstimulation of the cerebellum in monkeys showing evoked blood oxygenation level-depended (BOLD) fMRI activity in the dorsal pulvinar (Sultan et al., 2012). Indirect evidence for a functional connection could also be derived from a recently described human patient with a circumscribed (bilateral) lesion in the medial pulvinar who exhibited a mixture of parietal and cerebellar neurological deficits (Wilke et al., 2018). Being somewhat speculative, the anatomical and functional connectivity between pulvinar and cerebellum in primates needs to be studied in more depth.

### Methodological limitations and future directions

In our experiments, as in many previous studies, the head was immobilized in the straight ahead direction relative to the body and the center of the screen. Hence, head-, body- or world (screen)-centered references cannot be dissociated in this design (Mullette-Gillman et al., 2005). Owing to the same limitation, when interpreting the modulation of responses during fixation at initial or final gaze positions, we cannot distinguish between encoding of eye position in the orbit (e.g. via a proprioceptive signal) vs. spatial representation of the fixated stimulus (in any of the above reference frames).

In our design only 7 out of 15 final gaze positions resulted from different initial gaze positions and hence different saccade vectors, limiting the statistical power for dissociating the post-saccadic vector encoding from effects of post-saccadic eye position. Furthermore, we only varied the initial horizontal gaze position. A hypothetical extreme example of a purely retinocentric encoding with post-saccadic responses for all upper targets would also result in an apparent upper final gaze preference with the same ‘goodness of fit’– demonstrating that in some cases those two encoding schemes cannot be dissociated. Therefore, at least partially the final gaze encoding could be a result of “tiling” retinocentric (saccade vector) response fields originating from the three initial gaze positions. Several lines of evidence in our data however suggest that the static position of the eyes in the orbit is the defining factors in post-saccadic tuning, especially in the later part of the post-saccadic epoch. First, the corresponding target hold epoch analyzed here started 400 ms after the saccade end. Second, the two-way [initial gaze × retinocentric target position] ANOVA results on firing rates in target hold indicate that purely retinocentric encoding was rare: out of 156 units with the effect of the final gaze, only 27 units showed ‘purely retinocentric’ effect in the separate two-way ANOVA, while most showed also the effect of initial gaze and/or the interaction. Third, only 25 out of these 156 units showed an interaction between the initial gaze (i.e. saccade vector) and the final gaze in the two-way ANOVA on 7 positions that were reached from more than one initial gaze position (cf. Figure 4B). Fourth, an independent analysis that directly compared retinocentric vs. “screen-centered” (i.e. final gaze position-dependent) reference frames confirmed that most units were classified as the latter in the target hold epoch, despite relying on only a small number of overlapping target locations for calculating screen-centered correlations. Hence, the most parsimonious explanation of our results suggests that the eye position signals are indeed amply represented in the dorsal pulvinar.

It is important to note that the effects of initial vs. final gaze position were largely congruent (with a caveat of more extensive range for the final positions), and the influence of the preceding large saccade during the late part of the post-saccadic fixation was small. This implies that during initial fixation and target hold the cognitive task-contingent aspects were not a major factor in gaze-dependent modulation. We did not explicitly test for potential influences of small corrective saccades or preparation of return saccades after the end of the trial, but we did not observe any systematic relationship between patterns of the final gaze position dependence and retinocentric visual/saccade tuning in those units that showed such tuning.

In respect to encoding of cue, delay and saccade responses to the peripheral targets, our design allows dissociating between: i) purely retinocentric encoding, ii) retinocentric encoding modulated by gaze-dependent gain field, iii) head/body/world-centered, and iv) mixed reference frame. But it is important to note that with the current design (largely non-overlapping sets of final gaze positions linked to each of only three initial gaze position, as compared e.g. to 9 starting positions in (Andersen et al., 1990), or 9 horizontal targets in (Mullette-Gillman et al., 2005)), if a unit shows not a “straightforward” gaze-specific gain effect on retinocentric encoding with but a more complex pattern (e.g. shift or other idiosyncratic pattern), we cannot always reliably identify the best fitting reference frame (Mullette-Gillman et al., 2009). Future experiments with a larger range of initial and final target positions, and rotation of the head relative to the body, and the body relative to the screen (Dean and Platt, 2006), will elucidate the functional role of these signals in postural control and guiding visually-guided actions in space (Grieve et al., 2000).

Another potential limitation was that our experiments were not performed in total darkness, due to background illumination of the monitor. This could be considered as a potential confound because if a neuron has a large peripheral receptive field, the initial (or final) gaze position might bring this RF outside of the illuminated monitor. This is however unlikely to explain our findings for several reasons. First, the horizontal size of the screen was large (100 degrees of visual angle). Second, a roughly equal number of units showed same or opposite hemifield preference for visual cue and initial gaze position. If the effects of gaze were due to changes in visual stimulation, we would expect these preferences to be opposite (regardless of whether a unit is showing enhancement or suppression to a bright cue) because a larger part of the illuminated screen is always in the hemifield opposite to the direction of the gaze. Third, many units did not have a clear visual response but still showed the effect of gaze position. Finally, the two studies that tested the eye position effects in central thalamus and ventral pulvinar did not see a difference between light and darkness (Schlag-Rey and Schlag, 1984; Robinson et al., 1990). To summarize, the eye position dependence during fixation is most likely due to extraretinal signals or non-retinocentric spatial encoding.

### Functional significance of eye position signals in dorsal pulvinar

Generally, the functional significance of eye position signals in dPul is likely to be similar to other cortical and subcortical brain regions to which it connects, with the special property that the pulvinar is in a good anatomical position to distribute this information throughout the interconnected cortical circuitry (Sherman and Guillery, 2002; Saalmann and Kastner, 2011). We briefly list plausible functions before considering some of them in more detail.

In respect to eye position dependence during fixation, the putative functions could be:

1. Eye movements to auditory stimuli (Yao and Peck, 1997; Cohen and Andersen, 2002)
2. Computation of ocular vergence angle to determine depth/distance of foveated objects (Rosenbluth and Allman, 2002)
3. Maintaining stable fixation and smooth pursuit (Ohtsuka et al., 1991; Krauzlis et al., 2017)

In respect to influence of eye position on visual responses to peripheral targets and activity around saccades:

4. Neural computation of spatial target localization as an intermediate step in the transformation between retinal location and egocentric location in respect to the body or limbs (Andersen et al., 1993; Pouget and Snyder, 2000)
5. Modulation of cortical activity with predictive shifts of visual attention associated with saccades (Duhamel et al., 1992; Hoffman and Subramaniam, 1995; Wurtz et al., 2011b, 2011a)
6. Maintaining perceptual stability across eye movements (Sommer and Wurtz, 2008; Mirpour and Bisley, 2015)
7. Recalibration of efference copy signals after a saccade (Wang et al., 2007; Xu et al., 2012)

It is conceivable that the eye position signals in dPul are relevant for actions in depth. It seems possible that dPul neurons are sensitive for depth of fixation similar to fixation neurons in area 7a (Sakata et al., 1980). Indeed, pulvinar lesions can lead to stereoacuity deficits (Takayama et al., 1994). Since pulvinar neurons are binocular and are sensitive to relative retinal disparity, these cells may contribute to binocular depth perception (Casanova et al., 1991; Wilke et al., 2009). Other deficits after dorsal pulvinar lesions in monkeys and humans, involving nystagmus and impaired smooth pursuit suggest that it might be also important for gaze stabilization (Ohtsuka et al., 1991; Wilke et al., 2010, 2018).

Previous electrophysiological and inactivation studies showed that the dorsal pulvinar is critical for guiding visuospatial attention (Petersen et al., 1985; Fiebelkorn et al., 2019). The fact that the preference for a certain steady gaze direction was also found in dPul neurons that do not have a visual cue response suggests that the eye position signals are not only used to further modulate peripheral visuospatial attention. Instead, the modulation of firing during steady fixation might represent attention to relevant spatial targets, encoded either foveally or in a non-retinocentric reference frame. Many neurons decrease their firing during memory delay, as compared to initial fixation, suggesting that attention to peripheral locations and motor preparation might interact with foveal responses. Further work is required to dissociate these more cognitive possibilities from extraretinal orbital eye position signals.

While the functional role of gaze dependence in dPul can at least in part be derived from related properties in connected cortical areas and the superior colliculus, it should be constrained by known behavioral consequences of lesion and perturbation studies, as well as electrophysiological properties of dPul neurons. Given relatively coarse spatial tuning of most dorsal pulvinar responses to visual cues and saccades, and moderate effects of perturbation on saccadic precision and accuracy (Petersen et al., 1985; Robinson et al., 1986; Bender and Baizer, 1990; Benevento and Port, 1995; Wilke et al., 2010; Dominguez-Vargas et al., 2017), it is unlikely that the main function of eye position signals in dPul is precise perceptual or saccadic localization, as might be the case for SC and FEF (Caruso et al., 2018). The basic visuomotor characteristics of dPul seem to be closer to parietal areas LIP and 7a, as well as posterior cingulate (Andersen et al., 1990; Barash et al., 1991; Dean et al., 2004). There were however some differences between prevalent eye position effects in dPul and those cortical areas. In the pre-movement epochs, the encoding in dPul was mostly retinocentric or mixed, with a moderate occurrence of gaze-dependent gain fields. This is unlike more frequent gain fields in LIP (Andersen et al., 1990), or the large number of neurons with non-retinocentric encoding that has been cautiously interpreted as head-centered in parietal cortex (Mullette-Gillman et al., 2005; Caruso et al., 2018), or predominance of neurons exhibiting world-centered encoding in the posterior cingulate cortex (Dean and Platt, 2006). Nonetheless, the diverse patterns of eye position dependence in dPul can support at least some aspects of spatial transformation from retinocentric encoding to other coordinate frames during visuomotor planning.

The transient effects of saccadic displacement from one stable eye position to another seem to be of less relevance in the dPul given that the number of units showing the effect of the current eye position on pre-saccadic and peri-saccadic responses was small. Generally, some dPul neurons increase their firing predictively during the memory delay and/or just before the saccade and are tuned to specific retinocentric location, and might hypothetically participate in the transmission of the corollary discharge of the upcoming movement to the cortex, from the intermediate/deep layers of the superior colliculus or from one cortical area to another (Sherman and Guillery, 2002; Wurtz et al., 2011b). More neurons show pre/peri-saccadic decrease of firing, consistent with a contribution to saccadic suppression (Wurtz et al., 2011b). The effect of the eye position on the immediate post-saccadic responses is stronger, and this epoch shows a mixture between retinocentric encoding of the preceding saccade vector (with or without modulation by the pre-saccadic eye position), and the encoding of the current (post-saccadic) eye position. Post-saccadic eye position-related gain fields have been shown to provide unreliable localization signal contaminated by the pre-saccadic eye position, and thus their functional role should be interpreted with caution (Xu et al., 2012). A more detailed time-resolved analysis of the dynamics of eye position effects and reference frame transitions in the course of visual, memory delay, pre-saccadic and post-saccadic epochs is needed to elucidate the potential role of dPul eye position signals in preparation of the movements and post-movement updating.

Going beyond the oculomotor scope of this study, there are strong indications that the dPul is important for visually-guided reach and grasp movements (Acuna et al., 1990; Wilke et al., 2010, 2018), and likely eye-hand coordination (Grieve et al., 2000). In particular, the perturbation effects are not only space-specific, but also limb-specific, with deficits stronger for the contralesional hand (Wilke et al., 2010, 2018), and the dPul neurons respond differently when reaches are made with the left or the right hand (unpublished observations, see also (Domínguez-Vargas, 2017)). In the context of reach and grasp movements, the integration of eye position is a crucial step in spatial transformations from retinocentric to trunk-, limb-, or world-centered reference frames (Battaglia-Mayer et al., 2003; Batista et al., 2007; Marzocchi et al., 2008; Chang and Snyder, 2010; McGuire and Sabes, 2011; Bosco et al., 2015). In addition to limb movement aspects, gaze position has also been identified as an essential variable for stabilizing posture during upright standing (Ustinova and Perkins, 2011). Although this variable has not been explicitly addressed in monkey research, we have recently reported a patient with selective bilateral medial pulvinar lesions with difficulties to stand upright and walk, as well as nystagmus and hypometric saccades, without primary motor, vestibular or sensory deficits (Wilke et al., 2017, 2018). It is thus conceivable that the dorsal pulvinar contribution to integration of postural signals goes beyond oculomotor and reach/grasp movements, a possibility that should be explored in neurophysiological experiments.

## Supplemental Data

Supplemental data are available at https://figshare.com/s/846a1444f5f15cace27f

### Grants

Supported by the Hermann and Lilly Schilling Foundation, German Research Foundation (DFG) grants WI 4046/1-1 and Research Unit GA1475-B4, KA 3726/2-1, CNMPB Primate Platform, and funding from the Cognitive Neuroscience Laboratory.

### Disclosures

No conflicts of interest, financial or otherwise, are declared by the authors.

## Author contributions

L.S., A.U.D.V., M.W. and I.K. designed the experiments, A.U.D.V. and L.G. collected the data and performed initial analyses, L.S. analyzed the data, L.S. and I.K. prepared figures, L.S., M.W. and I.K. wrote the paper.

## Acknowledgements

We thank Ira Panolias, Daniela Lazzarini, Sina Plümer, Klaus Heisig, and Dirk Prüße for technical support. We also thank Stefan Treue, Alexander Gail, Hansjörg Scherberger, members of the Decision and Awareness Group, Sensorimotor Group and the Cognitive Neuroscience Laboratory for helpful discussions.

