## Supplemental Data for "Eye position signals in the dorsal pulvinar during fixation and goal-directed saccades"

3 Supplemental Figures

1 Supplemental Table

List of Abbreviations

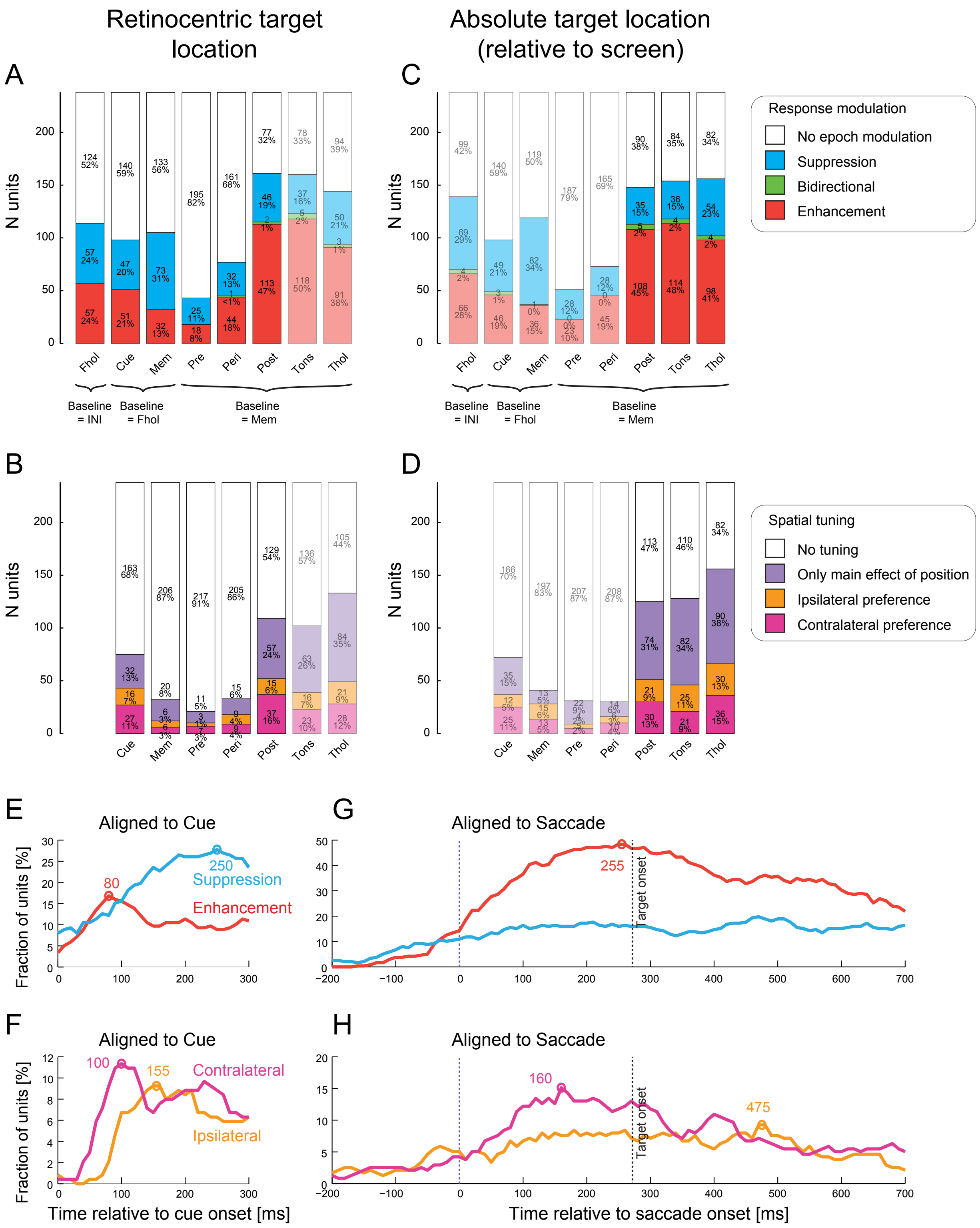


**Supplemental Figure S1. Spatial tuning and response modulation per epoch.** Response modulation and spatial tuning of all 238 units with more than 4 trials for each combination of initial gaze and retinocentric target location. The analyzed epochs are fixation hold (Fhol), cue onset (Cue), memory (MemE), pre-saccadic (Pre), peri-saccadic (Peri), post-saccadic (Post), target onset (Tons), and target hold (Thol). **A:** Total number of units (and percentages) showing response modulation relative to the respective baseline, for each analyzed epochs: no enhancement nor suppression (white), only suppression (blue), bidirectional - enhancement for one hemifield and suppression for the other (green), only enhancement (red). Target locations relative to the initial gaze position (retinocentric). **B:** Number of units that, in the respective epoch, were not tuned (white), did not prefer either hemifield but showed a main effect of position in a one-way ANOVA (purple), preferred ipsilateral hemifield (orange), preferred contralateral hemifield (magenta). Target locations relative to the initial gaze position (retinocentric). **C:** Same as A, but with target locations relative to the head/body midline or screen center (absolute target position on the screen). **D:** Same as B, but with target locations relative to the head/body midline or screen center.

**E-F:** Fraction of units showing modulation and spatial tuning for each 10 ms bin, relative to cue onset. Colored circles and numbers show the time points corresponding to the maximum number of modulated or tuned units. In case there were several maxima with the same amount of units, the respective time points were averaged. Note that a unit can contribute to multiple bins. **G-H:** Same as E-F, relative to saccade onset. Vertical lines show the saccade onset (purple) and the onset of the visible target (dotted lines). In all analyses, data across all initial gaze positions were combined. In E-H, target locations are relative to the initial gaze position (retinocentric). Note that epochs that are largely irrelevant for a given target reference frame are made semitransparent.

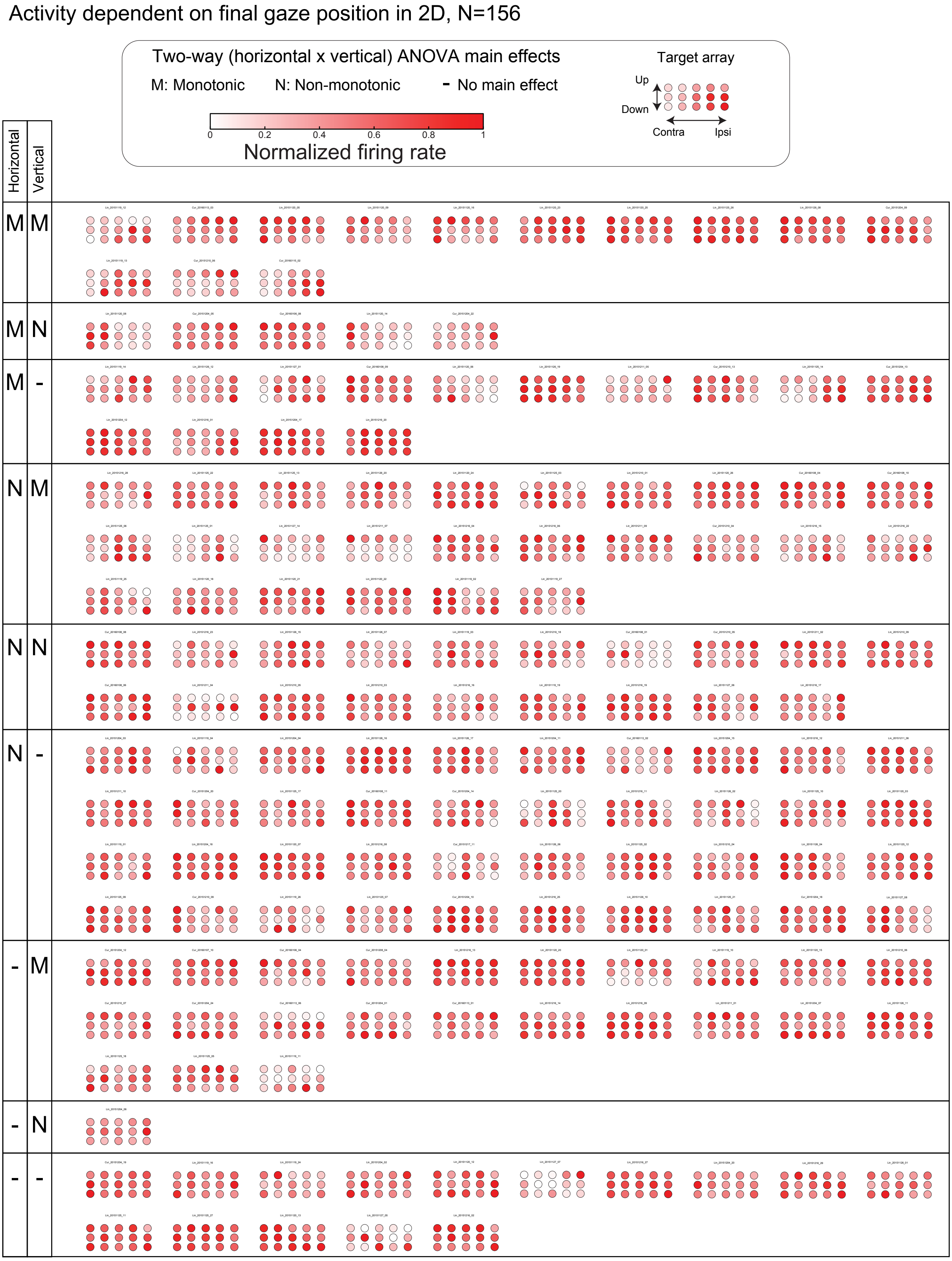


**Supplemental Figure S2. Final gaze tuning.** Heat map plots of firing rates for all 156 units that showed an ANOVA effect of final gaze position, normalized to the maximum firing rate. Units are grouped by monotonicity of vertical and horizontal gaze tuning: Monotonic (M), non-monotonic (N), and no main effect in the respective direction (-). Main effects of horizontal and vertical gaze position are derived from two-way ANOVA results.

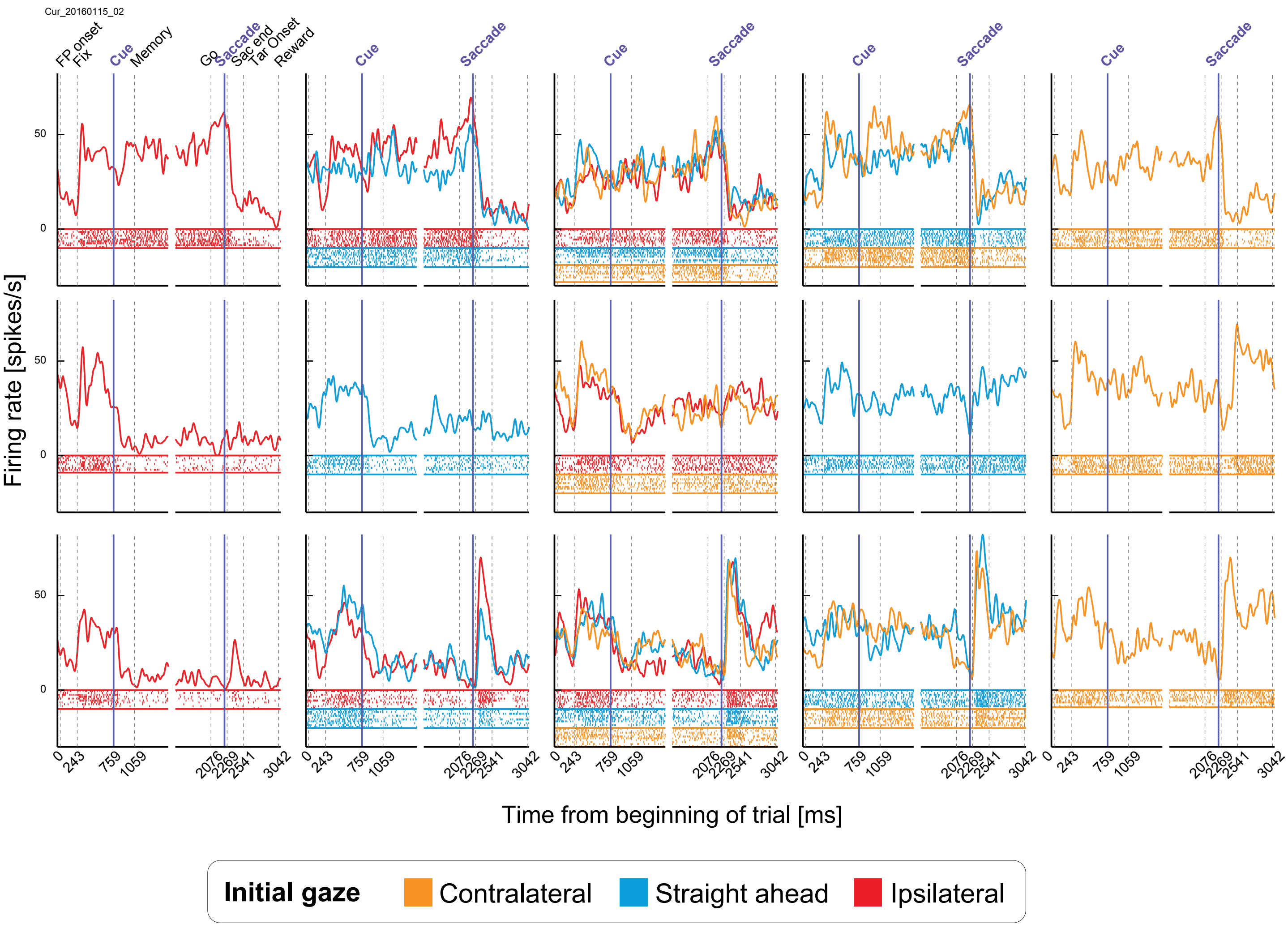


**Supplemental Figure 3. Example unit with mixed reference frame.** Example unit raster plots, spike density functions, and horizontal eye traces separately for each initial fixation position, for each of the 15 target locations relative to the screen. Colors depict the three initial gaze positions (orange: contralateral, blue: central, red: ipsilateral). This unit had high ACCs for both retinocentric and "screen-centered" references frames, from cue to pre-saccadic epoch, resulting in mixed reference frame classification.

| Fixation hold | | Pre-/Peri-/Post-saccadic response (retinocentric) | | | | |
| --- | --- | --- | --- | --- | --- | --- |
| | | Enhancement | Suppression | Hemifield preference | ANOVA position effect | No saccade response |
| Initial gaze effect present | 120 | 62 | 35 | 46 | 64 | 13 |
| Initial gaze effect absent | 118 | 64 | 35 | 45 | 58 | 15 |

**Supplemental Table S1. Relationship between initial gaze effect and saccade responses.** Number of units showing enhancement, suppression, hemifield preference, or an effect of retinocentric position during any of the three saccade-related epochs (pre-/peri-/post-saccadic), and none of the above, separately for units showing an initial gaze effect (in fixation hold, 120 units, cf. **Figure 4B,** 120=91+29) and units showing no effect of initial gaze (118 units).

### List of Abbreviations

| | |
|---|---|
| ANOVA | Analysis of variance |
| ACC | Average correlation coefficient |
| CI | Contralaterality index |
| LED | Light-emitting diode |
| MATLAB | Matrix laboratory |
| MRI | Magnetic resonance imaging |
| PEEK | Polyetheretherketone |
| RARE | Rapid acquisition with relaxation enhancement |
| PSTH | Peri-stimulus time histogram |
| SD | Standard deviation |
| **Anatomical** | |
| CL | Centrolateral nucleus |
| CM | Centromedian nucleus |
| dlPFC | Dorsolateral prefrontal cortex |
| FEF | Frontal eye field |
| IML | Internal medullary lamina |
| IPul | Inferior pulvinar |
| LGN | Lateral geniculate nucleus |
| LIP | Lateral intraparietal cortex |
| LPul | Lateral pulvinar |
| LP | Lateral posterior nucleus |
| MD | Mediodorsal nucelus |
| MIP | Medial intraparietal cortex |
| MPul | Medial pulvinar |
| MT | Medial temporal area |
| MST | Medial superior temporal area |
| PC | Paracentral nucleus |
| PLdm | Dorsomedial lateral pulvinar |
| PLvl | Ventrolateral lateral pulvinar |
| SC | Superior colliculus |
| TPO | Temporo-parieto-occipital area |
| VA | Ventral anterior nucleus |
| VIP | Ventral intraparietal cortex |
| VL | Ventral lateral nucleus |
